# Exploring New Horizons: A Novel Cdk5 Inhibitor Restoring Cognitive Function and Alleviating Type 2 Diabetes

**DOI:** 10.1101/2024.09.30.615976

**Authors:** Sangita Paul, Chandran Remya, K.V. Dileep, Juhi Bhardwaj, Praveen Singh, S Poornima, C Srinivas, A.M. Sajith, BK Binukumar

## Abstract

Type 2 diabetes (T2D) is a metabolic disorder frequently associated with cognitive decline, making T2D patients susceptible to dementia. Often referred to as type 3 diabetes, Alzheimer’s disease (AD) shares a close association with hyperglycemia and insulin dysregulation. Despite this, anti-diabetic medications have proven beneficial in reducing cognitive impairment induced by T2D. Previous research, including our own, has highlighted the dysregulation of Cdk5 activity in both T2D and AD, with downstream consequences contributing to the progression of pathophysiological changes in both disorders. Therefore, targeting the kinase Cdk5 may offer a more effective approach to treating T2D and cognitive deterioration. In our study, we present evidence supporting Cdk5 as a significant mediator between T2D and cognitive decline. Through the screening of the KINACore library, we identified novel brain-penetrant Cdk5 inhibitors, BLINK11 and BLINK15. Our study further validated the efficacy of these inhibitors in a high-fat diet-induced T2D model, demonstrating their rescue effects on T2D pathogenesis, including blood glucose levels, obesity, and cognitive impairment as assessed through behavioral studies. Notably, BLINK11 emerges as a promising Cdk5 inhibitor for improving the T2D phenotype and addressing cognitive impairment in T2D conditions.

**Graphical abstract:** 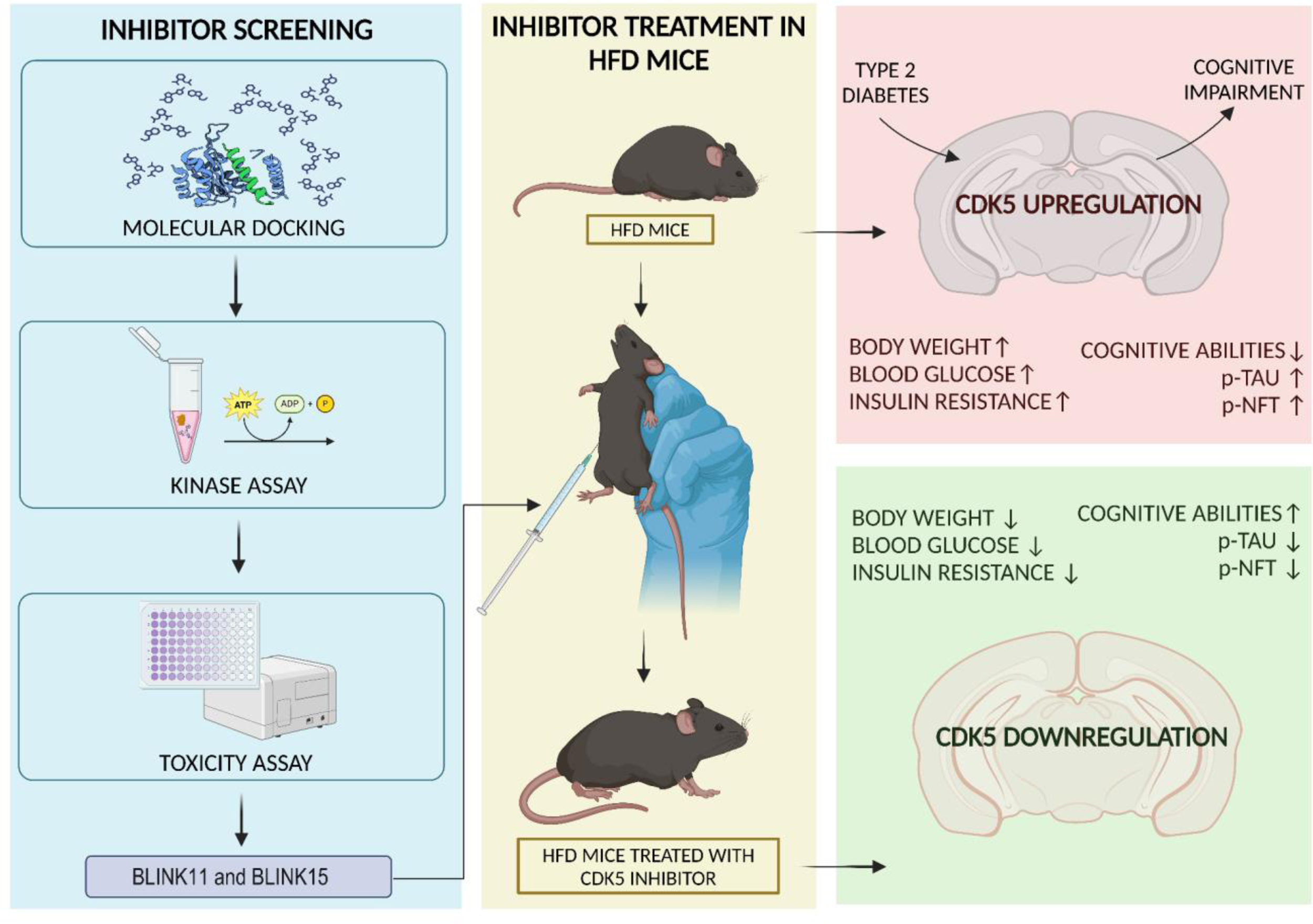

## Introduction

Diabetes is a long-term metabolic illness, and patients with type 2 diabetes (T2D) have been shown to experience a wide range of neurological abnormalities, including memory loss and cognitive decline (Li *et al*, 2023; Ciudin & Simó, 2022). Epidemiologically it has been proven that type 2 diabetic patients have a 70% increased risk of developing Alzheimer’s-like phenotypes such as neuroinflammation and cognitive impairment (Nakabeppu & Ninomiya, 2019; Butterfield *et al*, 2014; Hamzé *et al*, 2022). Insulin dysregulation has negative impacts on both peripheral glucose regulation and brain function since insulin is essential for both sustaining normal brain function and peripheral glucose metabolism. Numerous studies have indicated that an estimated two-to-three-fold relative risk for AD is associated with insulin resistance, which is characterized by unsuccessful glucose utilization (Chen & Zhong, 2013). This relationship may be explained by several variables, including oxidative stress, inflammation, mitochondrial dysfunction, and insulin-degrading enzyme (IDE) activity. Cyclin-dependent kinase 5 (Cdk5) can be a potent link between T2D and cognitive impairment as this kinase plays a critical role in both disease types (Lalioti *et al*, 2009; Liu *et al*, 2016). Studies have shown inhibition of hyperactive Cdk5 mitigates intracerebral neurotoxicity in diabetes and attenuates neuronal death and cognitive impairments (Liu *et al*, 2023; Chen *et al*, 2015).

Cdk5 is an atypical cyclin-dependent kinase that gets activated mainly through p35 and p39, predominantly in neuronal cells (Dhavan & Tsai, 2001). Its related downstream signaling pathways are known to strongly participate in several neurological events regulating neurodevelopment as well as neurodegeneration and cognitive impairment in the brain (Cheung & Ip, 2012; Paul *et al*, 2022; Bk *et al*, 2019). Cdk5 contributes to neurodevelopment and neuronal migration by phosphorylating various microtubules, actin-associated proteins, and cell motility proteins (Chae *et al*, 1997; Patrick *et al*, 1999). Again, cellular stress causes calcium influx into the cells enhancing calpain to cleave p35 into a truncated activator protein p25 which can more stably bind to Cdk5 and causes its hyperactivation. Contrastingly, p25 alters the substrate specificity of Cdk5, and (Patrick *et al*, 1999) favors its contribution to neurotoxicity. Partial knockdown of Cdk5 or inhibition of Cdk5 activity is considered a therapeutic strategy to rescue neurological disorders (Chae *et al*, 1997; Gutiérrez-Vargas *et al*, 2015; Patrick *et al*, 1999).

Interestingly studies have also shown the presence of Cdk5 and its activator p35 in pancreatic beta cells. The presence of glucose increases the expression of the p35 gene, which in turn encourages the creation of active Cdk5/p35 complexes controlling the expression of the insulin gene (Ubeda *et al*, 2004, 2006). A prolonged increase in glucose causes hyperactivation of Cdk5 which results in the inadequate release of insulin by pancreatic beta cells (Ubeda *et al*, 2006)). Therefore, dysregulation of Cdk5 is reported in T2D conditions and Cdk5 inhibition also shows antidiabetic effects (Liu *et al*, 2023). However, screening a novel Cdk5 inhibitor that efficiently inhibits Cdk5 can help rescue cognitive impairment induced by T2D in diabetic models.

## Results

### HTVS combined with MD simulations identified putative Cdk5 inhibitors

The compounds in the KINACore Library were designed based on two criteria: (1) molecules that share high structural similarity with known kinase activators and (2) molecules that have high structural similarities with the adenosine portion of ATP. The critical analysis of all Cdk5 structures revealed multiple conformations in the proximity of the active site, especially on the residues like E8, I10, K89, N144 (Fig 1 A). As all the structures are ligand-bound, we assumed that these conformational variations might exert an influence on ligand binding. Similarly, the receptor structures that are selected for our modeling studies include: (1) Cdk5-p25 in complex with indirubin (PDB ID: 1UNH), referred as R1, Cdk5-p25 in complex with roscovitine (PDB ID: 1UNL), referred as R2, Cdk5-p25 in complex with an ATP analog (PDB ID: 3O0G), referred as R3 and Cdk5 in complex with compound 4a (PDB ID: 4AU8), referred as R4. The ligands name in table 1 are indicated as L1-indirubin; L2-roscovitine; L3-{4-amino-2-[(4-chlorophenyl)amino]-1,3-thiazol-5-yl}(3-nitrophenyl)methanone; L4-4-(1,3-benzothiazol-2-yl)thiophene-2-sulfonamide (Table 1).

**Figure 1:**
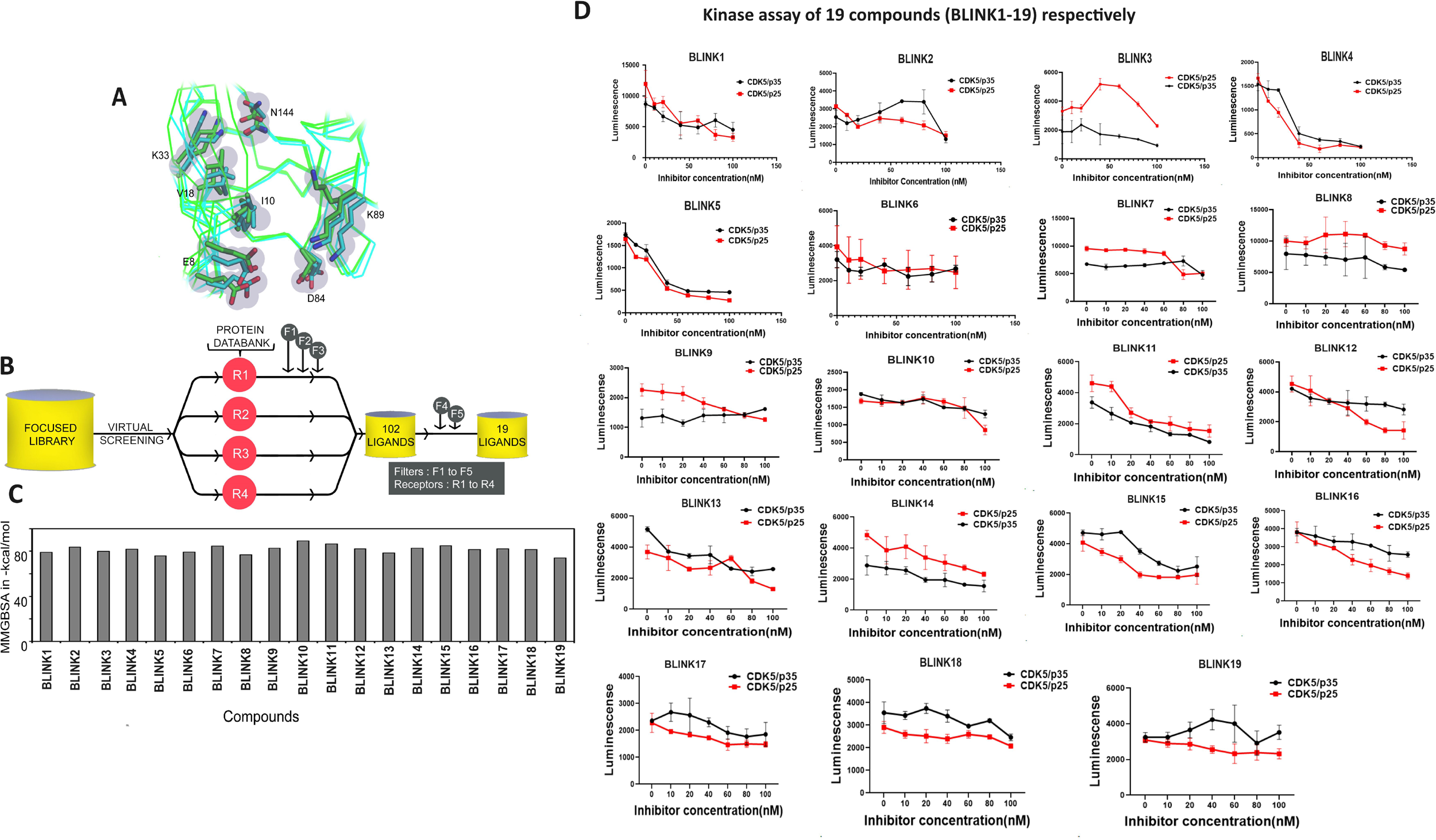
Virtual screening of Cdk5 inhibitors. (A) Superposition of different Cdk5 structures. The residues displaying distinct conformations are depicted in a stick model, while the protein backbone is illustrated using a ribbon model. Overview of the virtual screening workflow. (B) The ligands from the KINACore Library were screened against four distinct protein structures that differ in the orientation of the sidechains: PDB IDs 1UNH, 1UNL, 3O0G, and 4AU8, denoted as R1 to R4. These four receptors were employed for high-throughput virtual screening. The screening process comprised multiple filters, namely F1 to F5. F1, F2, and F3 corresponded to diverse docking protocols, specifically HTVS, SP, and XP docking, respectively. Additionally, F4 and F5 incorporated MM-GBSA calculations and visual inspections, respectively. (C) The binding energies of the final 19 ligands (BLINK1 to BLINK19) were determined using the MM-GBSA method. (D) Cdk5 activity in vitro with inhibitors in increasing concentration. Graphs 1-19 represent Cdk5 assay with p35 and 25 as activators with six different concentrations of BLINK1-19 (10-100 nM). IC50 was calculated for individual inhibitors with Cdk5/p35 and Cdk5/p25 complexes respectively. All the assays were performed in triplicates and 3 sets of data were considered for the analysis purpose.

**Table 1:**
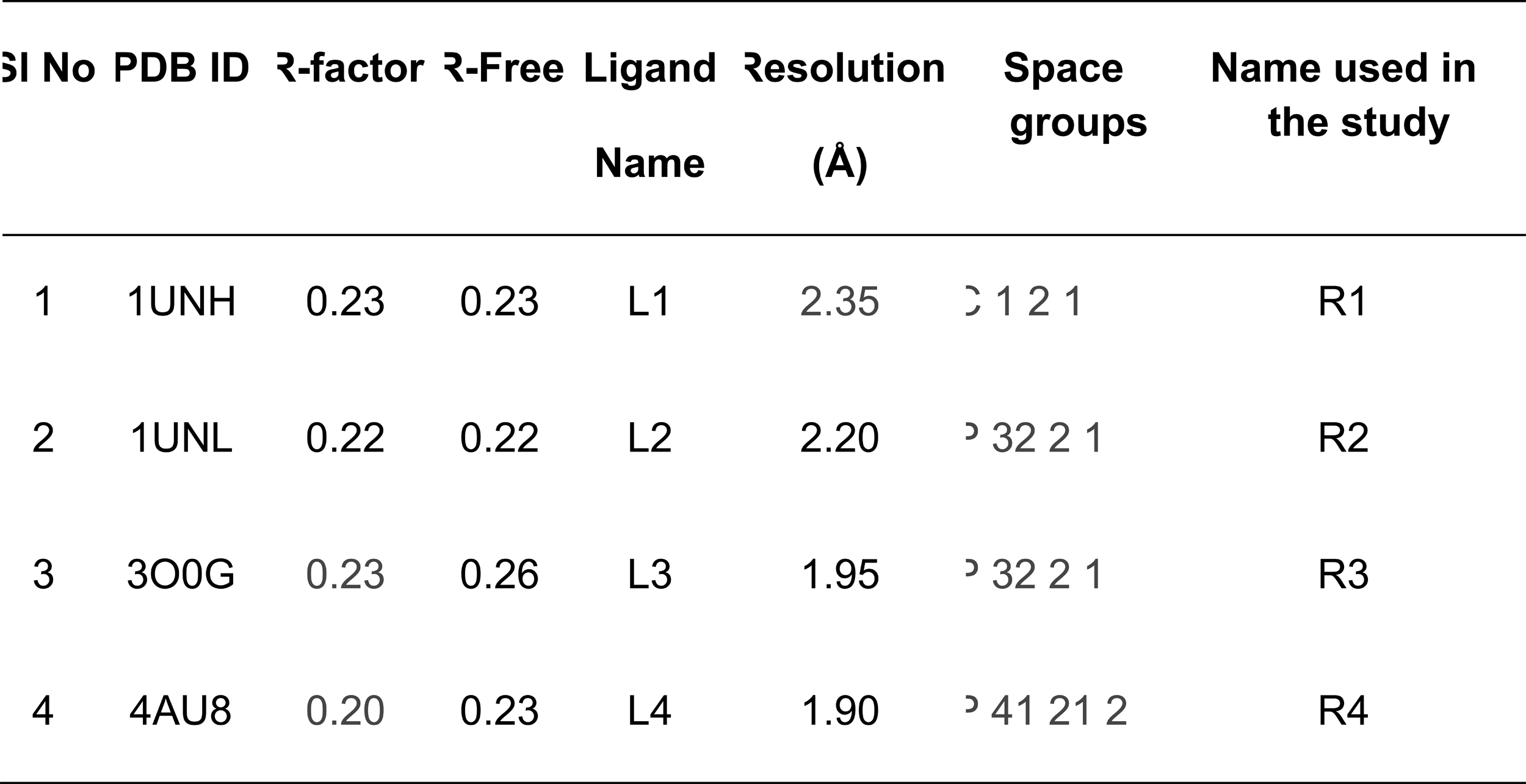
Overview of the Cdk5 crystal structures employed in the screening of KINACore libraries.

Our HTVS towards four receptor conformations identified 50 promising compounds for further studies (overall workflow of the in-silico studies shown in Fig1B). Interestingly, we observed that the screening against R1 and R2 is more energetically favorable. It appears that the conformation of I10 in R3 and R4 is apparently affecting the binding of ligands, reflected in lower binding energy. Finally, out of the 50 ligands, we selected 19 ligands (Table EV3) for the MD simulations. All these 19 ligands were named, BLINK1 to BLINK19 respectively, binding energies of these ligands are shown in Fig 1C. The structure of all the compounds is depicted in Fig EV2.

The MD simulation studies suggested that the ligands displayed versatile action on the active site of Cdk5 (Fig 2D and 2E and Fig EV3 A-Q).

**Figure 2:**
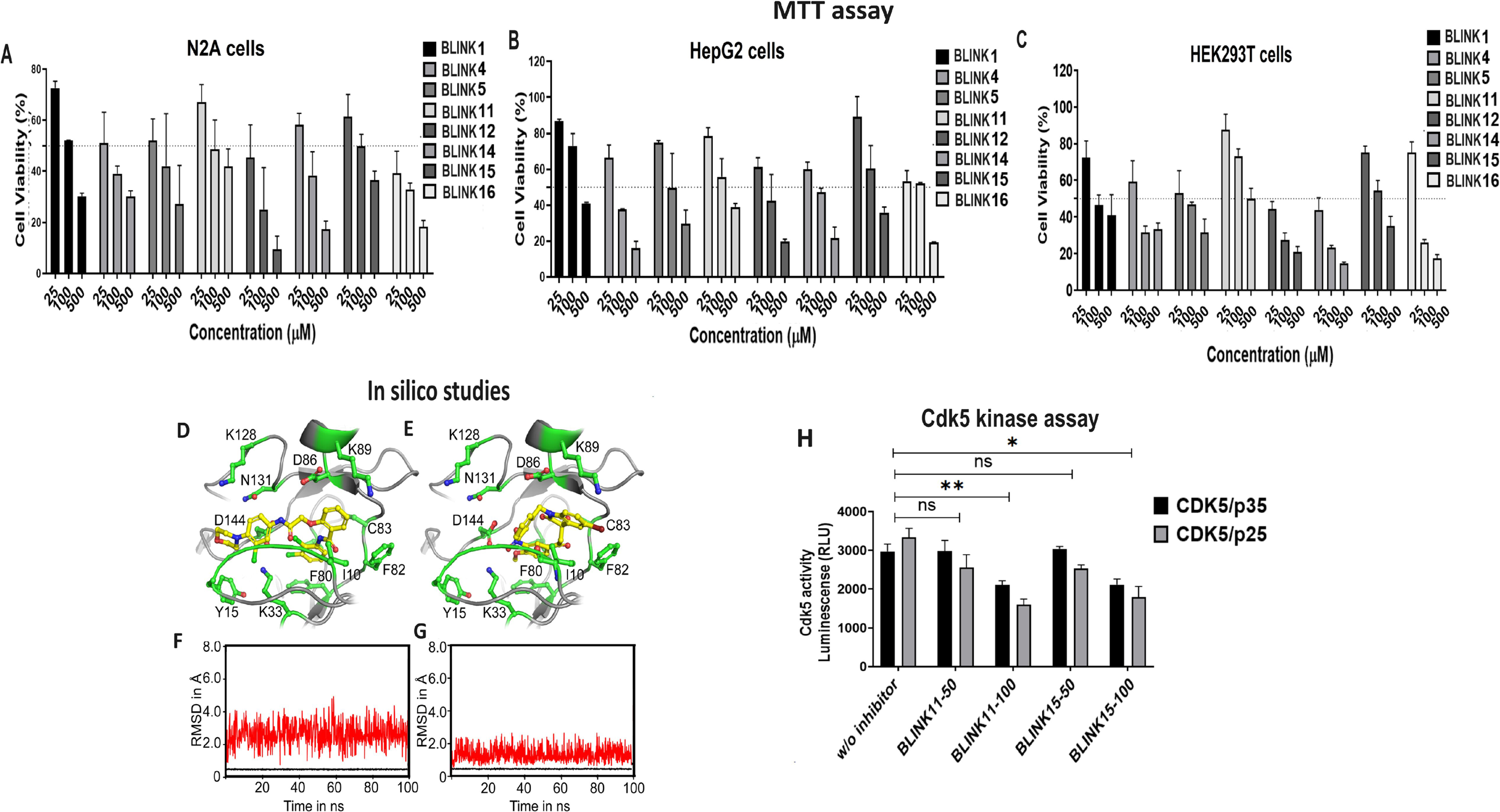
The cell viability test results for various inhibitors across different cell lines. The bar graph illustrates the percentage of cell viability, as determined by the MTT assay, for compounds BLINK1, BLINK4, BLINK5, BLINK11, BLINK12, BLINK14, BLINK15, and BLINK16 at concentrations of 25, 100, and 500 µM in (A) N2A cells, (B) HepG2 cells, and (C) HEK293T cells. All experiments were conducted in triplicates, and the data are presented as the mean ± standard deviation (SD). This analysis provides insights into the impact of these compounds on cell viability across different cell lines at varying concentrations. (D) Illustration of Ligand Binding Modes within the ATP Binding Pocket of Cdk5: the orientations of BLINK11, and (E) BLINK15 within the ATP binding pocket of Cdk5. The RMSD for BLINK11 (F), and BLINK15 (G) at the ATP binding pocket of Cdk5 over the course of a 100 ns simulation. The protein and the ligand RMSDs are shown in black and red lines respectively (H) The bar graph shows Cdk5 kinase activity in pulled down Cdk5 from cells overexpressed with Cdk5/p35 and Cdk5/p25 followed by BLINK11, and BLINK15 drug treatment at concentrations 50nM and 100nM

### In-vitro evaluation of putative Cdk5 inhibitors: Kinase assay, IC50 determination, and cell toxicity screening

The putative Cdk5 inhibitors were evaluated through an in-vitro screening process using the Cdk5 kinase assay. The luminescence observed was directly proportional to Cdk5 activity. Fig 1D presents the kinase assay results for all 19 inhibitors at increasing concentrations (0-100nM) for both Cdk5/p35 and Cdk5/p25 complexes. The calculated IC50 values are summarized in Table 2. Among the inhibitors tested, BLINK1, 4, 5, 11, 12, and 14-16 exhibited a more pronounced decrease in Cdk5 activity, suggesting their potential as effective inhibitors. These eight compounds were subjected to further screening based on cell toxicity using the MTT assay. The MTT assay was conducted on three different cell lines: HEK293T, HepG2, and N2A cells. The concentrations tested ranged from 25 to 100 uM (Fig 2A-C) indicating that compounds BLINK11, and BLINK15 maintained cell viability above 50% at all doses. This observation suggests a favorable safety profile for these compounds in terms of cell toxicity. The study highlights the importance of not only inhibiting Cdk5 activity but also ensuring minimal cytotoxicity, and these two compounds demonstrated promising characteristics in both aspects. We, however, proceeded our study with the two compounds BLINK11, and BLINK15, as they exhibited a more pronounced decrease in Cdk5/p25 activity when compared to other BLINKS. BLNK11 and BLNK15 also exhibit relatively lower activity against the Cdk5/p35 complex (see Table 2).

**Table 2:**
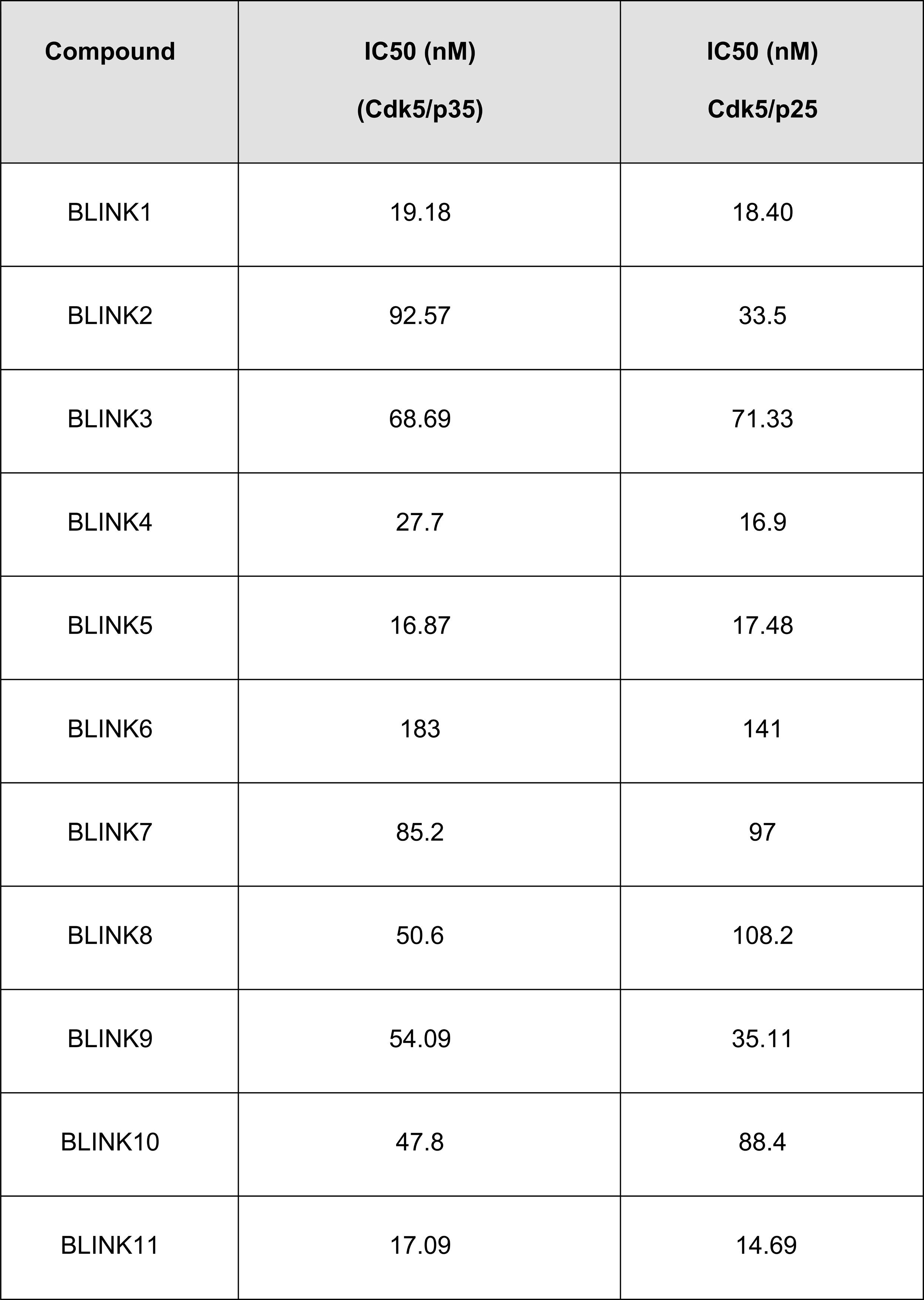
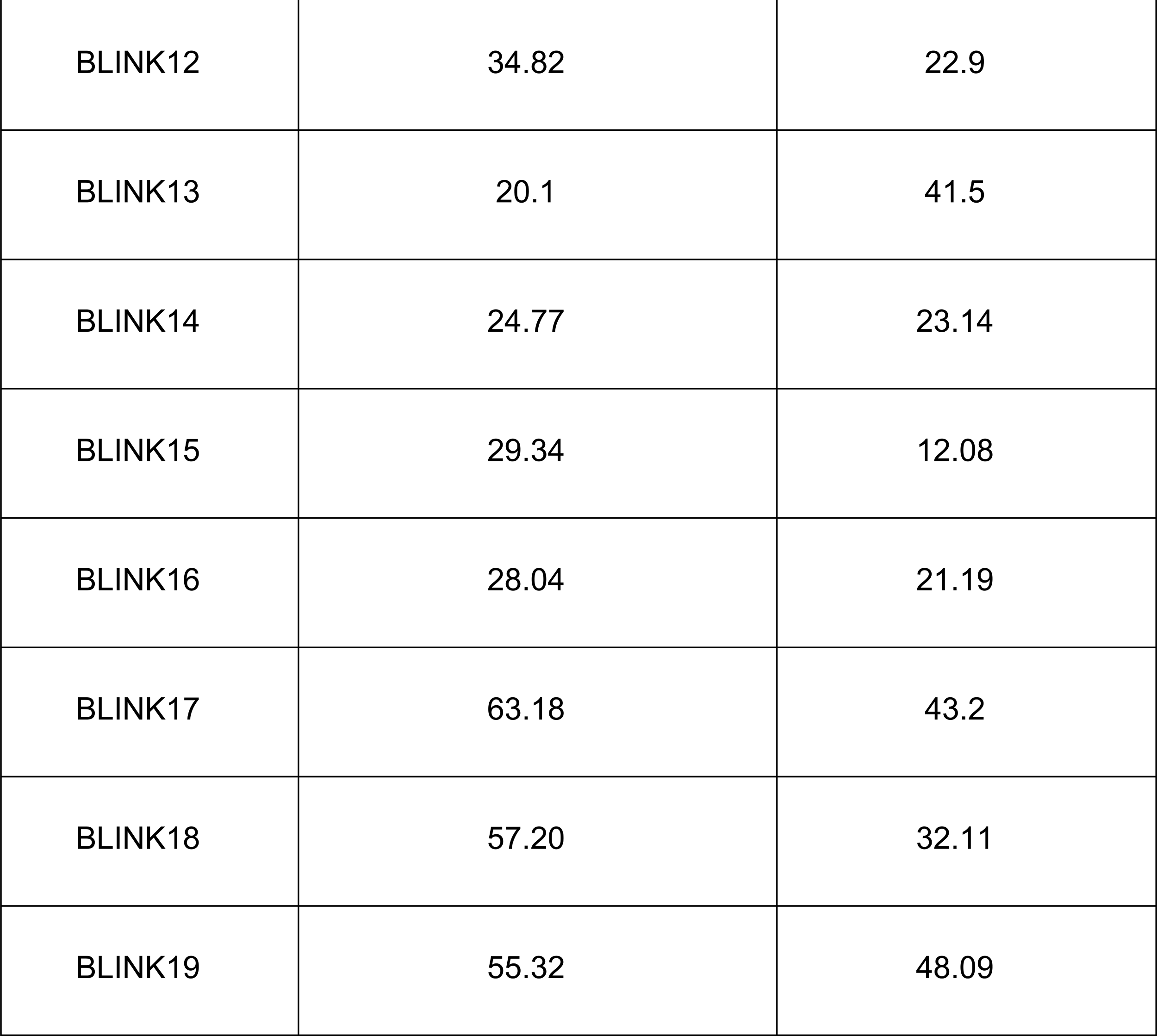
The IC50 calculated based on the kinase assay of Cdk5/p35 and Cdk5p25 complexes with all the inhibitors.

### Enhanced in vitro Activity of BLINK11 and BLINK15: Mechanistic Insights from In Silico Studies

The enhanced activity of BLINK11 and BLINK15 was explained through computational approaches. Molecular docking analysis revealed specific interactions of these ligands within the ATP binding pocket of Cdk5 (Fig 2D and 2E). The CDK5 inhibitors typically form hydrogen bonds with E81 and C83, while they also engage in hydrophobic interactions with residues I10, Q85, and L133 (Kalra *et al*, 2017).

Initially, BLINK11 forms a hydrogen bond with Q130 via its amide group but shifts to form a bond with C83 during MD simulations. Its morpholine ring also bonds with D144 of the DFG motif and interacts with D86 for over 85% of the simulation. An intramolecular hydrogen bond between its keto and amine groups adds conformational constraints. The dichloro phenyl benzamide moiety occupies the hinge region, facilitating interactions with C83, while the rest extends toward the DFG motif of the activation loop in Cdk5. BLINK11 also makes several hydrophobic interactions, involving residues like I10, C83, F82, L133, F80, V64, A31, A143, and V18. Fig 4A shows BLINK11 binding within the ATP binding pocket, while Fig 2F displays the RMSD for BLINK11 and protein during the 100ns simulation (Mapelli *et al*, 2005; Xing *et al*, 2015).

The orientation of BLINK15 at the ATP binding pocket is shown in Fig 2G. BLINK15 has been found to exhibit a similar hydrogen bonding pattern with C83 as observed in the case of other potent inhibitors of Cdk5. The carbonyl oxygen of C83 accepts a H-atom from the - OH group while the amide group donates a H-atom to the keto group of BLINK15. The interactions formed with C83 are found to be extremely stable during the MD simulation attributed to the inhibitory potency of BLINK15. In addition, the nitro group makes a salt bridge with K33 for ∼41% of the simulation time. The binding of BLINK15 was additionally stabilized by various hydrophobic interactions, similar to those observed in the case of BLINK11 contributing to the high binding stability of BLINK15 at the binding site of Cdk5 (Fig 2G).

### Cellular Impact and Blood-Brain Barrier (BBB) Permeability Assessment of Cdk5 Inhibitors BLINK11, and BLINK15

Our in vitro and in silico studies clearly demonstrated the inhibitory effects of compounds BLINK11, and BLINK15 against CDK5. Expanding our research to a cellular context, we conducted experiments in N2A cell lines overexpressing both CDK5/p35 and CDK5/p25. After subjecting the cells to two different concentrations, namely 50nM and 100nM, of each compound for a 24-hour period, we observed a significant decrease in CDK5 activity, as illustrated in Figure 2E. This noteworthy reduction in CDK5 hyperactivity at the 100nM concentration of BLINK11, and BLINK15 within the N2A cell model underscores the potential utility of these inhibitors in modulating CDK5 activity in a cellular environment. Also we noticed a decreased activity for CDK5/p25 than CDK5/p35 in the presence of both BLINK11 and BLINK15 at 100nM treatment, emphasizing their relevance for further exploration in addressing conditions associated with CDK5 dysregulation.

Our next step involved evaluating the ability of these compounds to cross the BBB, a crucial consideration in drug targeting for the central nervous system. To address this challenge, we employed the PAMPA for BLINK11, and BLINK15. The results revealed that compounds BLINK11 and BLINK15 exhibited high permeability through the BBB, with Paap values exceeding 10 x 10-6 (Fig EV4 A-B). This elevated permeability suggests that these compounds possess the capability to effectively traverse the BBB and access the central nervous system. This finding supports their candidacy for addressing CDK5 hyperactivity in neurodegenerative conditions associated with T2D. The ability of BLINK11 and BLINK15 to efficiently cross the BBB enhances their potential as therapeutic agents for targeting CDK5-related mechanisms in the central nervous system.

To assess the specificity of inhibitors BLINK11 and BLINK15 against CDK5, we investigated their effects on different Cdks, i.e Cdk1, Cdk2, and Cdk9 that share structural homology with CDK5. Our findings indicate that Cdk1, in conjunction with its activator cyclin Y, exhibited inhibitory patterns with IC50 values of 39.34nM for BLINK11 and 35.84nM for BLINK15. However, these IC50 values were higher than those observed for CDK5/p35 and CDK5/p25. Conversely, Cdk2 (with cyclin A2) and Cdk9 (with cyclin T) showed minimal inhibition by both BLINK11 and BLINK15, with significantly higher IC50 values (Fig EV4 C-F). Overall, BLINK11 and BLINK15 can selectively target CDK5 at lower concentrations.

### LC-MS analysis reveals the bioavailability and distribution of BLINK11 and BLINK15 compounds in the HFD mouse model

To assess the BBB penetrability of BLINK11 and BLINK15 in the HFD mouse model, intraperitoneal injections were administered, and LC-MS analysis was conducted on both brain lysate and serum samples (Fig 3). The LC-MS results depicted in Figure 4.9A and B showcase the peaks corresponding to the BLINK11 compound in the brain lysate and serum, respectively. Similarly, Fig 3C and D illustrate the peaks associated with the BLINK15 compound in the brain lysate and serum, respectively. The bar graph in Fig 3E represents the area under the curve of the peaks for BLINK11 in both serum and brain, while Fig 3F depicts the corresponding values for BLINK15 in serum and brain. These analyses provide a quantitative measure of the presence of the compounds in the respective samples. Furthermore, to ascertain the relative distribution of BLINK11 and BLINK15 in the brain compared to serum, the percentages of each compound present in the brain were calculated and plotted in a bar graph (Figure 3G). This assessment helps to elucidate the ability of these compounds to traverse the blood-brain barrier and accumulate in the brain tissue. The LC-MS data presented in Figure 3 collectively contributes crucial insights into the bioavailability and distribution of compounds BLINK11 and BLINK15 in both the serum and brain lysate of the HFD mouse model. This information is pivotal for understanding the pharmacokinetics of the administered compounds and their potential impact on the central nervous system in the context of the studied neurodegenerative model.

**Figure 3:**
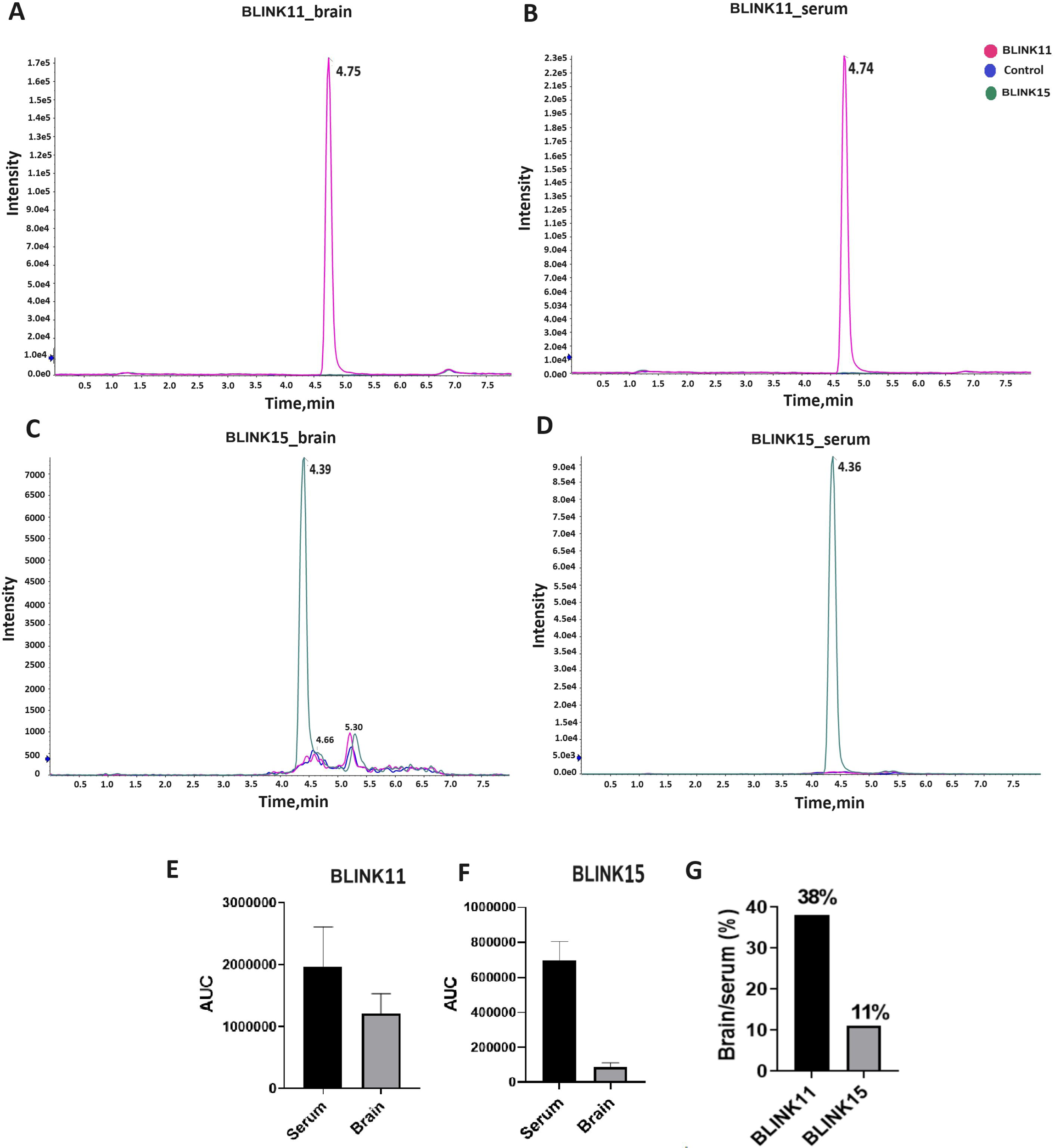
Bioavailability and distribution of compounds in brain lysate and serum. (A) The graph shows the peaks of the BLINK11 compound in the brain lysate, (B) serum (C). The graph shows the peaks of the BLINK15 compound in the brain lysate (D) serum The bar graph represents the area under the curve of the peaks of (E) BLINK11 in serum and brain (F) BLINK15 in serum and brain (G) Percentages of BLINK11 and BLINK15 present in brain compared to serum is calculated and plotted in a bar graph.

### Effect of Cdk5 inhibitor treatment on body weight, glucose tolerance, insulin tolerance, and plasma insulin levels in rescued T2D mice

The HFD-fed mice model received intraperitoneal injections of the compounds BLINK11 and BLINK15 at doses of 20 mg/kg/day and 40 mg/kg/day for a total of 20 days in the third month of HFD feeding. The body weight was measured at the beginning of the drug therapy, on day 0, and then again on days 10 and 20, as indicated in the Fig 4A. The findings demonstrate a fast and steady increase in body weight in the 3-month-old high-fat diet-induced mice group on days 10 and 20 compared to the control group following the administration of the drug (Fig 4A).

**Figure 4:**
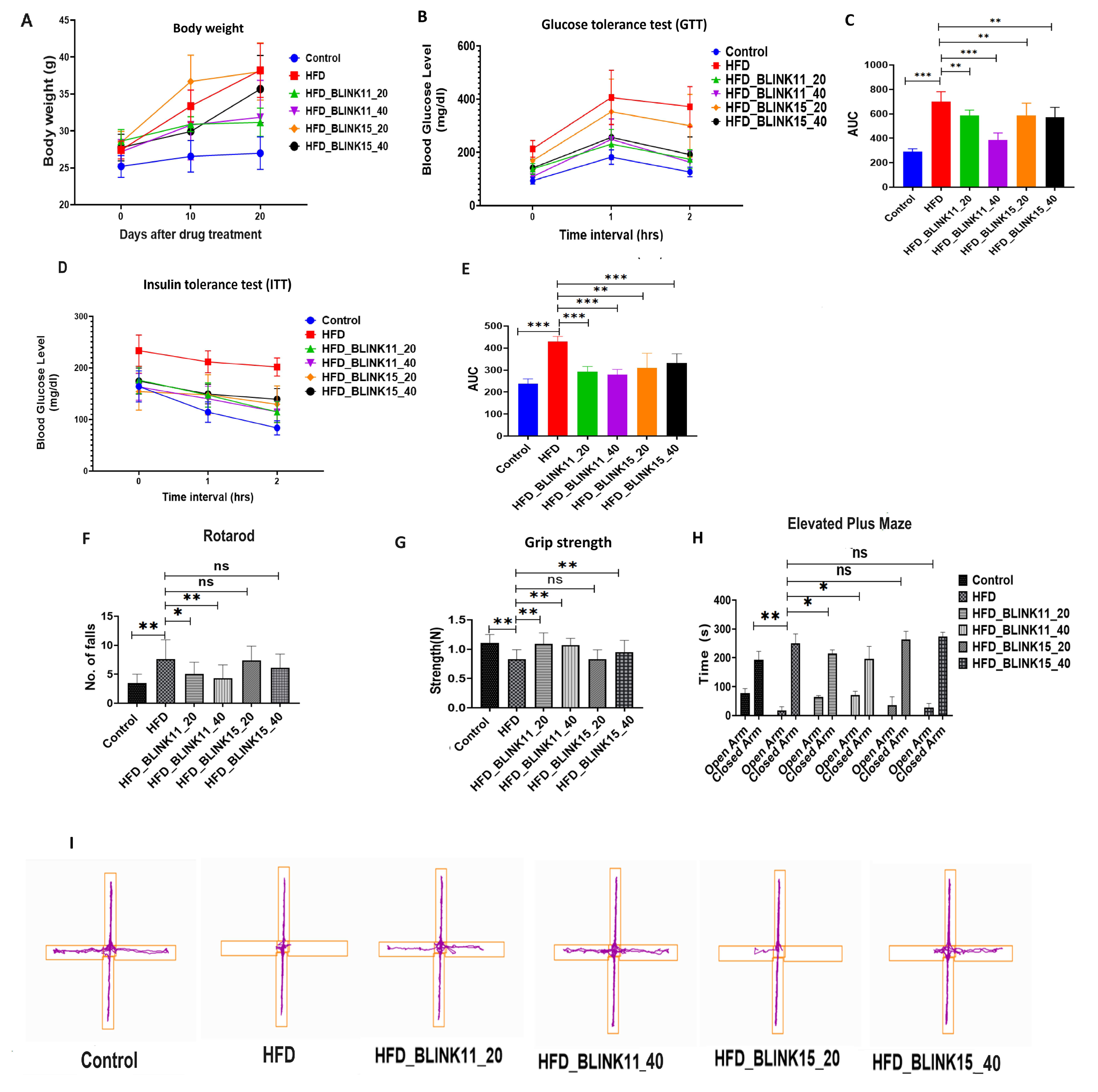
The establishment of a high-fat diet–induced diabetes mouse model with biochemical parameters is shown in graph. (A) The graph showing body weights of control, HFD, HFD treated with BLINK11 20mg/kg/day (HFD_BLINK11_20), HFD treated with BLINK11 40mg/kg/day (HFD_BLINK11_40), HFD treated with BLINK15 20mg/kg/day (HFD_BLINK15_20) and HFD treated with BLINK15 40mg/kg/day (HFD_BLINK15_40) at 0,10 and 20 days followed by drug treatment (n=8 per group) (B) Glucose tolerant test (GTT) (C) Area under the curve (AUC) plots for GTT per group. (D)Insulin tolerant test (ITT). (E) Area under the curve (AUC) plots for ITT per group. Tukey’s multiple comparison test and two-way paired ANOVA were used for the analysis, and the values were provided as mean± SD; *p>0.05, **p>0.01; ***p>0.001. **Behavior of mice in diverse experimental paradigms** (F) Graph showing the number of falls in different mice groups while performing the rotarod test (G) Grip strength measurements of the mouse groups. (H) Elevated Plus Maze results show time spent by the mice in open-arm and closed-arm. (I) Representative track plots in EPM of different mice groups. All the experiments were performed with n=8 per group. Tukey’s multiple comparison test and one-way ANOVA were used for the analysis, and the values were provided as mean± SD; *p>0.05, **p>0.01; ***p>0.001.

The BLINK11 drug-treated group is demonstrated to be rescued from the gain in body weight caused by a high-fat diet for both dosages; however, for the BLINK15 drug-treated group, only the higher dose, i.e. 40 mg/kg/day, exhibits significant recovery (Fig 4A). GTT and ITT were used to check the biochemical parameters for T2D in mouse models. Compared to the control group, the HFD group showed disturbed GTT and ITT, but the mouse group treated with BLINK11 and BLINK15 with both the dosages demonstrated a significant rescue in both GTT and ITT (Figure 4B-E).

### Effectiveness of Cdk5 inhibitors BLINK11 and BLINK15 in alleviating cognitive impairment: Insights from behavioral experiments in high-fat diet mouse models Cdk5 inhibitor treatment rescues muscle strength and alleviates anxiety in a T2D mouse model

Following the assessment of the peripheral antidiabetic protective effects of the compounds BLINK11 and BLINK15, our investigation extended to evaluating a battery of behavioral tests commonly impaired in the context of T2D. Initially, we focused on examining muscle strength and coordination using the rotarod and grip strength tests. The rotarod test revealed notable findings in terms of forelimb muscular strength among the various experimental groups. Specifically, mice subjected to a high-fat diet exhibited a significant decline in muscle strength compared to the control group. This observation suggests that a high-fat diet may adversely impact an animal’s ability to build muscle and motor coordination. Encouragingly, mice from the high-fat diet group treated with BLINK11 demonstrated a rescue effect, indicating a mitigation of the deterioration in muscle strength and motor coordination on the rotarod (Fig 4F).

In the grip strength test, the administration of compounds BLINK11 and BLINK15 at a dose of 40 mg/kg/day exhibited a rescue behavior, as illustrated in Figure 4G. This suggests that these specific treatments contributed to an improvement in grip strength compared to untreated counterparts. These findings indicate the potential therapeutic efficacy of BLINK11 and BLINK15 in addressing muscle strength deficits associated with T2D, emphasizing their role in mitigating the adverse effects of a high-fat diet on musculoskeletal function.

Elevated anxiety levels have been identified as another notable concern in subjects with T2D. To investigate anxiety behaviors among the mice groups, we conducted an elevated plus maze (EPM) test, focusing on the time spent by each mouse in the open and closed arms of the apparatus. The results of the EPM test revealed distinctive patterns in anxiety-related behaviors. Mice subjected to a high-fat diet displayed heightened anxiety compared to the control group, as evidenced by increased time spent in the closed arm of the maze Figure 4.11C, 4.11D). Intriguingly, treatment with BLINK11 in the high-fat diet group exhibited a rescuing effect on anxiety behavior (Figure 4H, 4I), indicating a potential amelioration of anxiety levels associated with T2D. Contrastingly, the group treated with

BLINK15 in the high-fat diet showed no significant difference compared to the high-fat diet group, as illustrated in Fig 4H and 4I. This suggests that while BLINK11 treatment demonstrates efficacy in alleviating anxiety behaviors, the effects may not be universally applicable across all compounds. These findings underscore the nuanced impact of specific treatments on anxiety-related behaviors in the context of T2D, warranting further exploration into the mechanisms underlying these observations.

### Rescue of memory and spatial learning impairments in T2D mouse models with Cdk5 inhibitor treatment

Memory and spatial learning are known to be impaired in the later stages of T2D. To investigate these impairments and explore potential drug treatments, we conducted Y-maze and water maze experiments on T2D mouse models. In the Y-maze, the alteration percentage of the HFD group was significantly lower than that of the control group, indicating memory and spatial learning deficits. Notably, mice in the HFD group treated with BLINK11 at a dose of 20 mg/kg/day did not exhibit any rescue effect, and similar results were observed with the BLINK15 compound treatment when compared to the untreated HFD group. However, in the HFD group treated with BLINK11 at a dose of 40 mg/kg/day, the decrease in alteration percentage was recovered. Additionally, the time spent in the altered arm showed no significant differences between the groups (Fig 5A, 5B).

**Figure 5:**
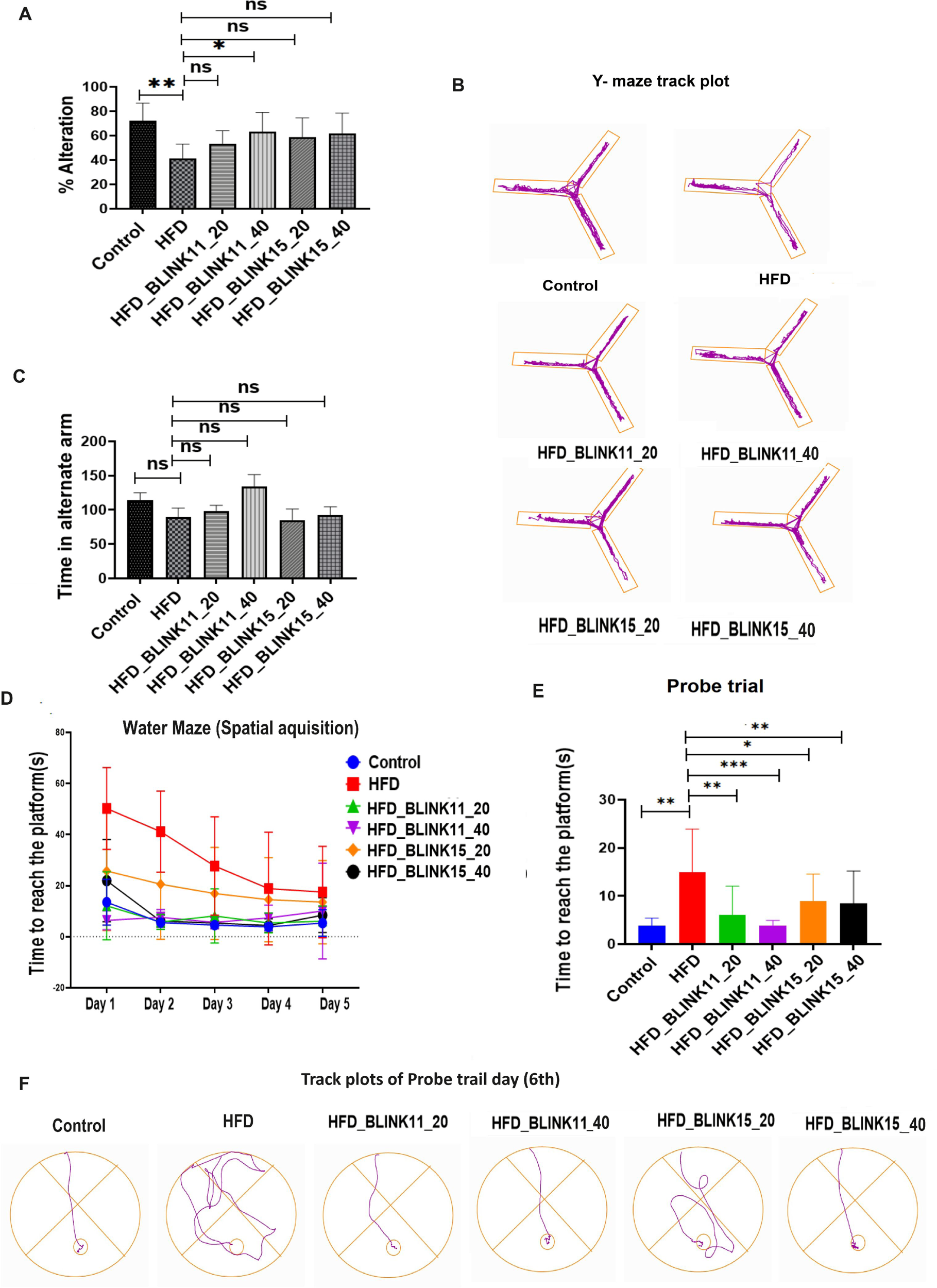
Behavioral study for memory and special learning through Y maze and Morri’s water maze. (A) The graph represents the percentage alteration within the mice groups Control vs HFD and HFD vs BLINK11, BLINK15 treated groups in Y maze (B) The images are the track plots of the mice groups in Y maze apparatus (C) The graph showing time spent by the mice in altered arms of the Y maze apparatus. (D) The graph plots the time taken by the mice of different groups to reach the platform in the water maze during the training period from Day 1 to Day 5 (E) The bar graph shows the time taken by the mice per group to reach the platform in the probe trial day (F) The images represent track plot of water maze performed by the mice groups in the probe trial day. All the experiments were performed with n=8 per group. Tukey’s multiple comparison test and one-way ANOVA were used for the analysis, and the values were provided as mean± SD; *p>0.05, **p>0.01; ***p>0.001.

Results from the water maze demonstrate a progressive learning pattern from the first to the fifth training days, with HFD mice models taking longer to reach the platform than the control group. In the HFD mouse treated with BLINK11 and BLINK15 at both doses, we can witness rescue behavior in this instance as well. According to the findings, the HFD mouse exhibits a reduction in spatial learning and memory. Treatment with BLINK11 and BLINK15 may be able to reverse this cognitive impairment brought on by a high-fat diet and obesity. Compared to the BLINK15 treatment, the BLINK11 treatment in HFD exhibits a more remarkable improvement (Fig 5D-F).

The results obtained from the water maze experiment unveiled a discernible pattern of progressive learning over the course of five training days. Notably, the HFD mouse models exhibited a prolonged duration to reach the platform compared to the control group, indicative of spatial learning challenges associated with HFD and obesity. However, in the HFD group treated with BLINK11 and BLINK15 at both doses, a noteworthy rescue behavior was observed. The cognitive impairment related to spatial learning and memory in HFD mice seemed to be mitigated with the administration of Cdk5 inhibitors BLINK11 and BLINK15. The data suggested that both compounds held the potential to reverse the detrimental effects induced by a high-fat diet. Interestingly, when comparing the two treatments, the BLINK11 treatment in the HFD group exhibited a more remarkable improvement in spatial learning and memory, as illustrated in Fig 5D-F. These findings underscore the therapeutic potential of Cdk5 inhibitors, particularly BLINK11, in ameliorating cognitive deficits associated with T2D conditions induced by an HFD, providing valuable insights for future research and potential clinical applications.

### Enhanced Cdk5 activity and p25 generation in the HFD model rescued by Cdk5 inhibitor treatment

In our investigation of the molecular pathways contributing to cognitive impairments in the T2D mouse model, we have previously documented an association between elevated CDK5 activity and increased p25 generation in the brains of HFD mice, correlating with cognitive deficits (Saha et al., 2023). This study focuses on the hippocampal lysate of HFD mice, revealing consistent findings. The presence of p25 leads to heightened CDK5 activity in the brains of HFD mouse models. Notably, treatment with CDK5 inhibitors BLINK11 and BLINK15 results in a significant reduction in CDK5 activity, even in the presence of p25. Intriguingly, the hippocampal expression levels of CDK5 remain unaltered following treatment. Additionally, p25 generation in the BLINK11 and BLINK15 treated groups remains unchanged, as depicted in Fig 6

**Figure 6:**
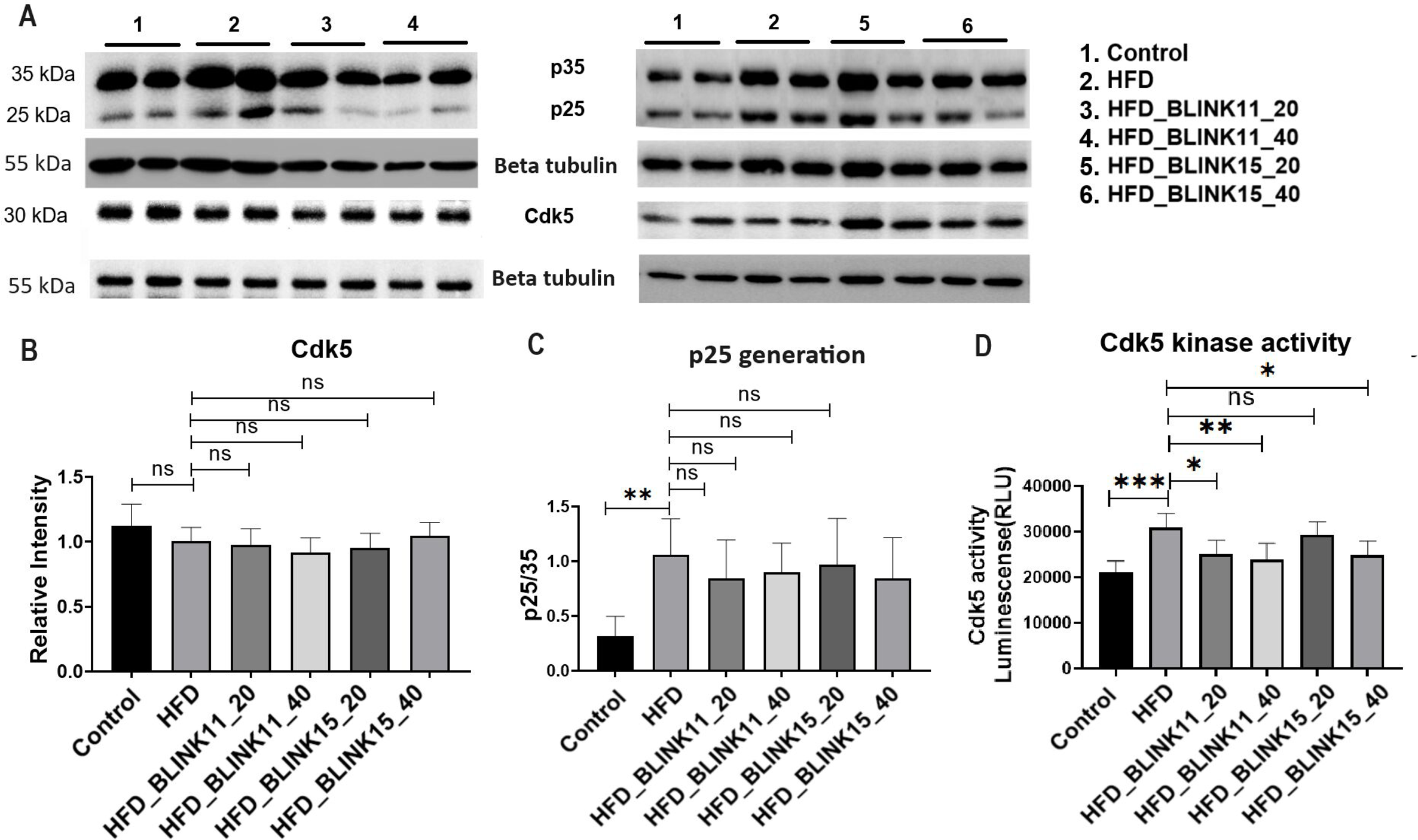
Cdk5 expression and activity along with p25 generation in treated vs non treated HFD mouse groups. (A) Western blot showing expression patterns of p35/25 and Cdk5 in the hippocampus region of mouse brain among the mouse groups control, HFD, and HFD treated group. (B) Densitometry of the respective blots Cdk5 with respect to the control beta-tubulin and p25 with respect to p35 intensity. (C) Cdk5 kinase activity using pulled down Cdk5 from the hippocampus lysate of the mouse groups. All the datasets were analyzed using one-way ANOVA and Tukey’s multiple comparison tests, the data were provided as mean±SD, n=6. **p<0.05, **p<0.01, ***p<0.001

### Enhanced expression patterns of neurodegeneration markers in the HFD model are alleviated by the treatment of Cdk5 inhibitor BLINK11

The heightened cognitive deficits in the HFD mouse model prompted an exploration into neurodegenerative markers within the hippocampus. Despite unchanged Neurofilament Heavy chain (NF-H) levels, phospho-NF-H was notably increased in the HFD group compared to the control. The Cdk5 inhibitor BLINK11, particularly at a higher dose, significantly reduced phospho-NF-H levels. Additionally, neurofilament light chain (NF-L) levels, indicative of neurodegeneration, were diminished in the HFD group but restored with BLINK11 treatment. Tau and phospho-tau (231Thr and 396Ser) levels were substantially elevated in the HFD group, but BLINK11 treatment significantly decreased phospho-tau-396Ser levels (Fig 7).

**Figure 7:**
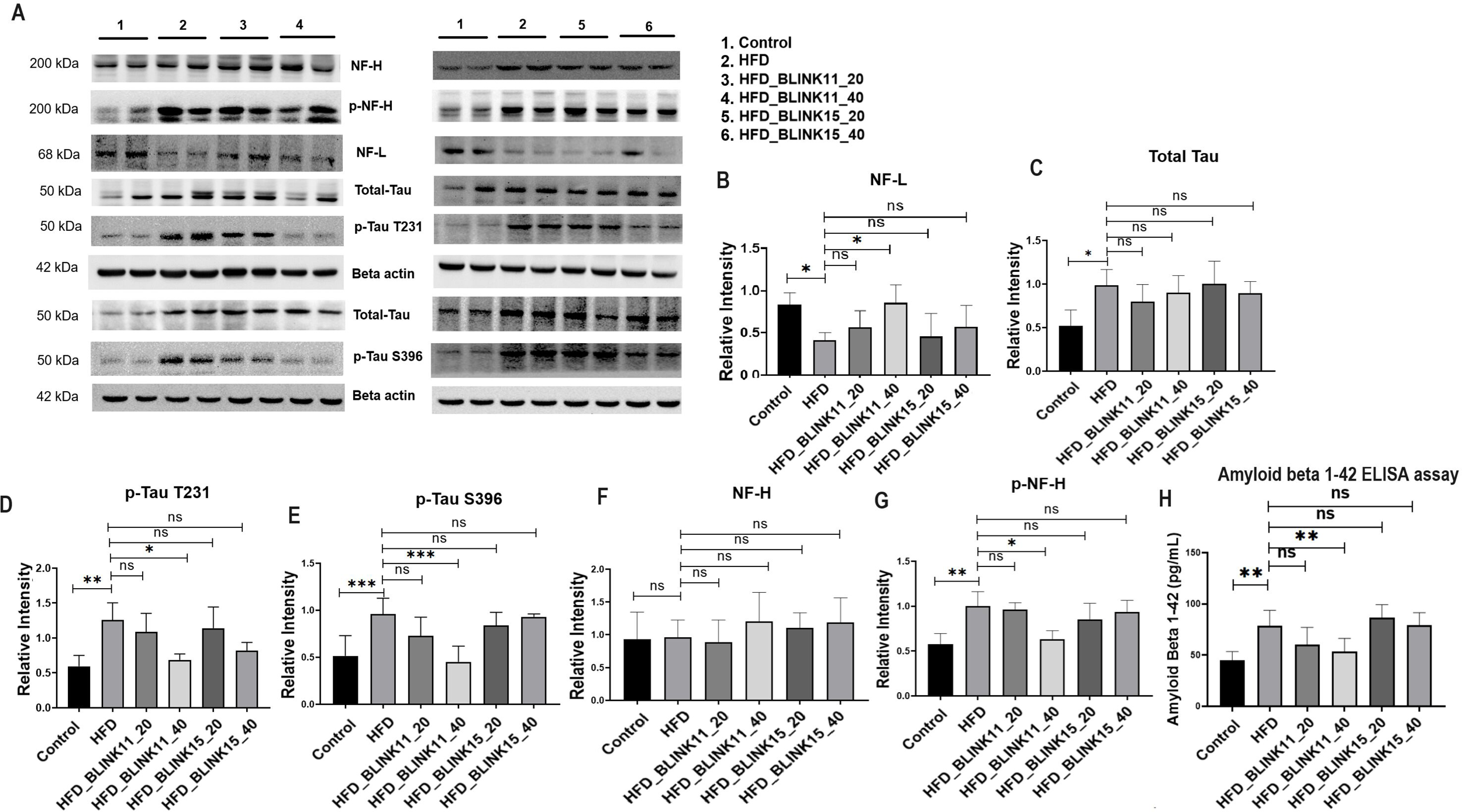
Expression patterns of neurodegenerative markers. (A) The western blot images to check the expression pattern of NF-H, p-NF-H, NF-L, Tau, p-Tau-T231, p-Tau-S396 in the hippocampus region of mouse brain among control, HFD non-treated and HFD treated groups. (B-G) The densitometry analysis of the respect blots NF-L, total tau, Tau, p-Tau-T231, p-Tau-S396, NF-H and p-NF-H. The ratio of the protein expression of p-tau and NF-H to the protein expression of total tau and NH-H was used to normalize the relative expressions. The expression levels of the rest of the proteins have been normalized with beta-actin. (H) Bar graph showing quantification of amyloid beta 1-42 in pg/ml in the mice groups using ELISA assay. Datasets were analyzed using one-way ANOVA and Tukey’s multiple comparison tests and were provided as mean±SD, n=6. **p<0.05, **p<0.01, ***p<0.001

Immunohistochemistry on hippocampal sections confirmed these findings, showing consistent expression patterns of NF-L, p-Tau 231Thr, p-tau 396Ser, and total Tau with the western blot results (Fig 8). These results suggest that Cdk5 inhibition by BLINK11 mitigates HFD-induced neurodegeneration through the inhibition of phosphorylation events involving NF-H and tau proteins. Moreover, ELISA assays revealed a significant increase in amyloid beta 1-42 (Ab1-42) in the HFD group, which was significantly reduced by BLINK11 at 40mg/kg. Thus, BLINK11 at this dosage effectively rescues neurodegeneration caused by HFD.

**Figure 8:**
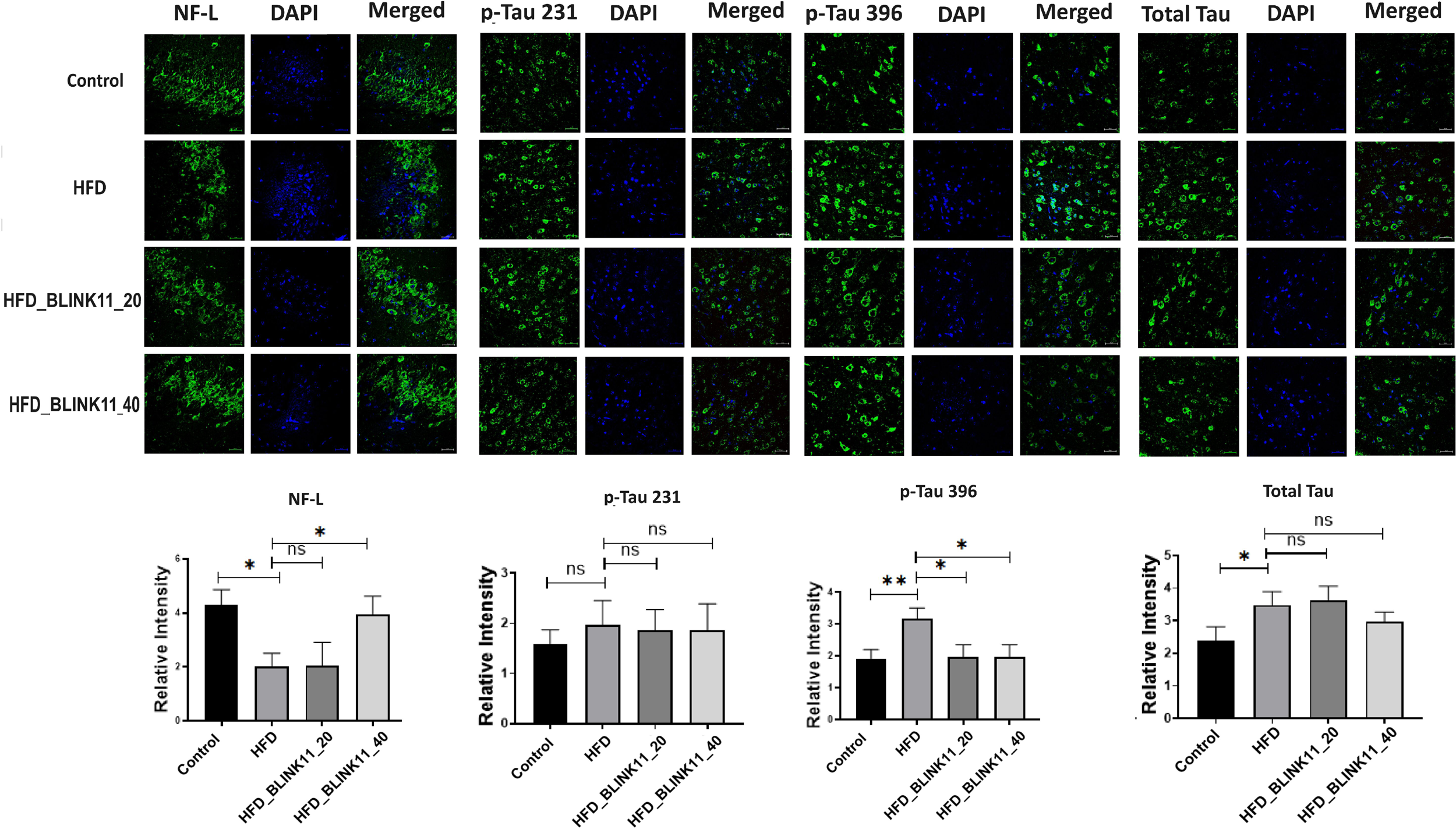
Immunohistochemistry results showing the expression patterns of. (A) NF-L, total Tau, p-Tau T232, p-Tau S396 in the hippocampus region of the brain sections sliced with 10-micron thickness. (B) Bar graph showing the intensity of the respective proteins NF-L, total Tau, p-Tau T232, and p-Tau S396 with respect to their corresponding DAPI intensities. All the datasets were analyzed using one-way ANOVA and Tukey’s multiple comparison tests, the data were provided as mean±SD, n=6. **p<0.05, **p<0.01, ***p<0.001

## Discussion

The global rise in the prevalence of T2D and AD is a growing concern, emphasizing the need for a deeper understanding of their pathophysiology (Rathmann & Giani, 2004; 2020 Alzheimer’s disease facts and figures, 2020). The strong evidence linking these two conditions underscores the importance of exploring shared mechanisms that contribute to their progression. Chronic glucose toxicity has been identified as a major contributor to neurological impairments associated with diabetes, providing a foundation for investigating potential therapeutic interventions (Rojas-Carranza *et al*, 2018; Patrone *et al*, 2014; Paul *et al*, 2024). CDK5 hyperactivation has emerged as a key player in the progression of neurodegeneration, neuroinflammation, and cognitive impairment in various neurological disorders (Paul *et al*, 2022; Ao *et al*, 2022; Binukumar *et al*, 2014, 2015b; Binukumar & Pant, 2016). Neurofilament tangles (NFT) production, protein aggregation, tau-phosphorylation, and neuroinflammation are influenced by dysregulation of CDK5 (Liu *et al*, 2016; Lu *et al*, 2019). Multiple studies, including the present one, have demonstrated that inhibiting CDK5 can significantly improve neurodegenerative phenotypes (Pao *et al*, 2023; Wang *et al*, 2020; Grant *et al*, 2017; Shukla *et al*, 2017). The study builds on prior research indicating that high glucose exposure leads to CDK5 hyperactivity and oxidative stress, resulting in hyperphosphorylation of tau (Ubeda *et al*, 2006; Zheng *et al*, 2013; Binukumar *et al*, 2014). Importantly, the current investigation extends these findings to suggest that CDK5 hyperactivation under glucotoxicity may contribute to the loss of beta cell function and insulin resistance, proposing CDK5 inhibitors as potential candidates for the treatment of T2D.

Previous research, including our work, has highlighted the hyperactivation of CDK5 and the generation of p25 in HFD mouse models of T2D, coinciding with cognitive decline (Zheng *et al*, 2013; Cao *et al*, 2022; Saha *et al*, 2023). This study takes a significant step forward by screening a novel library of CDK5 inhibitors for their potential to rescue T2D phenotypes, cognitive impairments, and neurodegeneration in a T2D mouse model induced by a high-fat diet.

The initial phase of the study involved a comprehensive in-silico screening of the KINACore library for CDK5 inhibitors using molecular docking. The library was meticulously designed based on structural similarities with known kinase activators and the adenosine portion of ATP. The identification of 50 promising compounds through high-throughput virtual screening (HTVS) laid the groundwork for further *in vitro* evaluations. The top eight compounds with low IC50 values were selected for additional screening, considering both their efficacy and low cytotoxicity across multiple cell lines (Fig 1 and 2).

To assess the BBB penetration and bioavailability of the selected compounds, LC-MS analysis was conducted on brain lysate and serum samples (Fig 3). The results indicated distinct peaks corresponding to the compounds BLINK11 and BLINK15, emphasizing their potential to reach the central nervous system. Subsequent evaluation of biochemical parameters relevant to T2D phenotypes in HFD mice treated with BLINK11 and BLINK15 revealed a rescue in weight gain, improved glucose tolerance, and reduced insulin levels compared to the untreated HFD group. These findings support the potential of CDK5 inhibitors in mitigating peripheral complications associated with T2D (Fig 4).

Moving beyond peripheral effects, the study explored the impact of CDK5 inhibitors on behavioral and cognitive aspects in the HFD mouse model. The assessment of muscle strength and coordination through the rotarod and grip strength tests demonstrated notable deficits in the HFD group, while treatment with BLINK11 exhibited a rescuing effect. Additionally, both BLINK11 and BLINK15 treatments showed a rescue behavior in the grip strength test. Elevated anxiety levels, commonly associated with T2D, were assessed using the elevated plus maze (EPM) test. BLINK11 treatment exhibited a rescuing effect on anxiety behavior, highlighting its potential in ameliorating anxiety levels associated with T2D (Fig 4F-I).

Cognitive impairments, a significant concern in both T2D and neurodegenerative diseases, were evaluated through Y-maze and water maze experiments. The results indicated significant deficits in memory and spatial learning in the HFD group, with BLINK11 treatment showing a potential cognitive benefit. Both BLINK11 and BLINK15 treatments demonstrated a rescue behavior in the water maze, with BLINK11 exhibiting a more remarkable improvement in spatial learning and memory (Fig 5).

The study delves into the molecular pathways underlying cognitive impairments in the T2D mouse model induced by an HFD. Elevated CDK5 activity and increased p25 generation in the brains of HFD mice were consistent with prior findings, and treatment with CDK5 inhibitors, particularly BLINK11, resulted in a significant reduction in CDK5 activity. Importantly, BLINK11 treatment effectively reversed elevated phospho-NF-H levels and restored diminished expressions of NF-L (Ashton *et al*, 2019; Hoyer-Kimura *et al*, 2021), indicating a potential neuroprotective effect. Tau hyperphosphorylation, a hallmark of neurodegeneration, was substantially reduced with BLINK11 treatment, further supporting its therapeutic potential. Immunohistochemistry results reinforced these findings, providing a comprehensive understanding of the impact of CDK5 inhibition on neurodegenerative markers (Fig 7A-G).

Quantification of amyloid beta 1-42 (Ab1-42) in the brain lysate revealed a significant increase in the HFD group, with BLINK11 treatment showing a significant rescue effect (Fig 7H). These results collectively underscore the therapeutic efficacy of CDK5 inhibition, particularly with the compound BLINK11, in alleviating neurodegeneration induced by an HFD in mouse models. The modulation of neurodegenerative markers, including NF-H, NF-L, and tau phosphorylation, highlights the potential of CDK5 inhibitors as promising therapeutic interventions for neurodegenerative conditions associated with metabolic disorders.

In conclusion, this study provides a comprehensive investigation into the potential of novel CDK5 inhibitors, particularly BLINK11, in addressing both peripheral and cognitive complications associated with T2D induced by an HFD. The multi-faceted approach, combining in-silico screening, *in vitro* evaluations, and *in vivo* assessments, strengthens the validity of the findings. The demonstrated rescue effects on muscle strength, anxiety levels, and cognitive impairments underscore the therapeutic potential of CDK5 inhibitors, offering promising avenues for further research and clinical applications. The study not only contributes to our understanding of the molecular mechanisms underlying T2D-associated neurodegeneration but also positions CDK5 inhibition as a compelling strategy for mitigating the complex interplay between metabolic disorders and cognitive decline.

## Materials and Methods

### Screening of Cdk5 inhibitors from KINACore

The KINACore library, containing over 20,000 drug-like molecules from ChemBridge, was screened for Cdk5 inhibitors. The library was prepared using the LigPrep module of Schrödinger (Schrödinger, LLC, New York, USA), generating different tautomeric and ionization states via the Epik program, followed by energy minimization using the OPLS-2005 force field. These prepared ligands were then subjected to high throughput virtual screening against Cdk5 (Shelley *et al*, 2007; Harder *et al*, 2016).

### Selection, preparation of Cdk5 structure, and high throughput virtual screening

Structural analysis of Cdk5 from PDB revealed conformational changes in amino acids upon ligand binding. Four distinct receptor conformations were selected for screening. These structures were prepared using Schrödinger Maestro’s Protein Preparation Wizard, with steps including the addition of polar hydrogen atoms, protonation states optimization and hydrogen bond network optimization. Energy minimization was performed with an OPLS-2005 force field. A grid around the bound ligands’ centroid was generated for docking studies. The virtual screening protocol involved HTVS, SP, and XP dockings, selecting the top 30% of compounds at each step for MM-GBSA calculations using the Schrödinger Prime module.

### Molecular dynamic simulation

To further observe the stability of the Cdk5-ligand complexes for a period of time, we performed molecular dynamics simulations (MD-Simulations) using Desmond software with GPU acceleration (Bowers *et al*). The Cdk5-ligand complexes with top binding energies were selected for MD-simulations. Using the System Builder module, we prepared the protein-ligand complex for MD simulation by keeping them in an orthorhombic simulation box with a dimension of 5 x 5 x 5 Å containing TIP3P water molecules. To neutralize the system, a required number of Na+ or Cl-ions was added. MD simulation was performed using an OPLS-2005 force field for 100 ns, with a recording interval of 100 ps. We utilized the NPT ensemble, maintaining constant temperature (Fixed at 300K) and pressure (fixed at 1.01 bar). Temperature and pressure control were implemented through the Nose-Hoover chain and Martyna-Tobias-Klein methods, respectively. Integration was performed by utilizing the RESPA algorithm with a time step of 2 fs and default configurations were applied to all other simulation parameters

### Compound synthesis

The 19 compounds screened through molecular docking are procured from Chembridge for initial in-vitro screening. The bulk synthesis was done for in-vivo experiments. The synthesis steps for BLINK11 are as follows Reactions were set-up on fume hoods and carried out under nitrogen atmosphere in Schlenk tubes unless otherwise noted. Compounds were purified by flash chromatography (Isolera-Biotage) using silica gel (200-400 mesh). All the reagents and solvents were used as received from commercial sources, unless otherwise specified. 1H NMR data was recorded on Bruker 400MHz AVANCE series or Bruker300 MHz DPX Spectrometer with CDCl3 or DMSO-d6 as solvent. 1H chemical shifts were referenced to CDCl3 at 7.26 ppm and for DMSO-d6 at 2.51 and 3.20 ppm. Multiplicities are abbreviated as follows: singlet (s), doublet (d), triplet (t), quartet (q), doublet-doublet (dd), quintet (quint), sextet (sextet), septet (septet), multiplet (m), and broad (br). MS was measured on Agilent 7890A/5975C Series GC/MSD mass spectrometer or Agilent 1100 Series LC/MSD mass spectrometer.

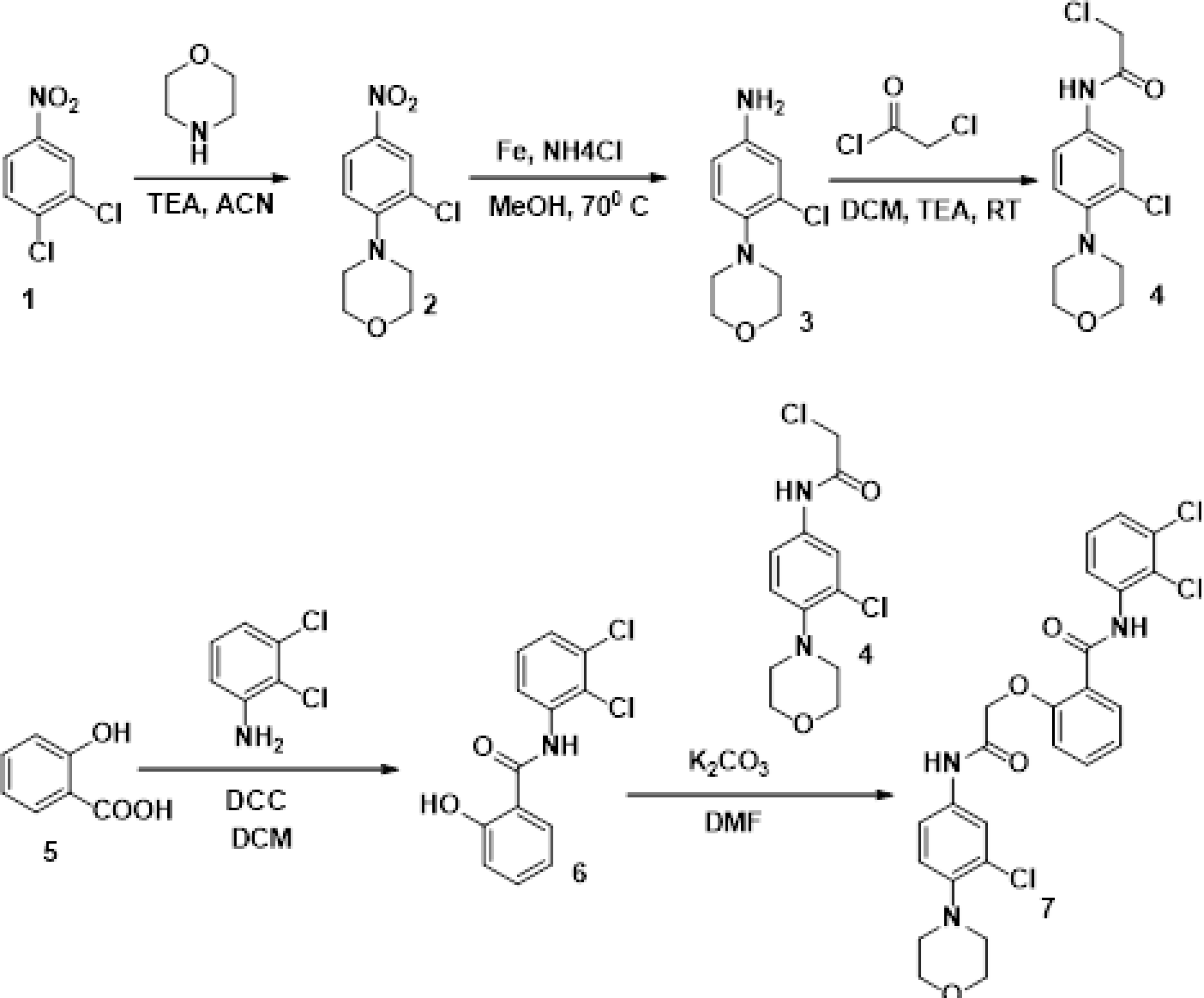

Scheme 1: Synthesis of compound 7

The synthesis of target compound 7 for biological evaluation involves the above synthetic sequence depicted in Scheme 1. The synthesis of intermediate 4 was first accomplished via 3 steps. The first step involves the nucleophilic displacement of 3,4 dichloro nitro benzene with morpholine, followed by reduction and subsequent reaction with chloro acetyl chloride to yield compound 4. The synthesis of compound 6 involves the amide formation reaction of compound 5 and 2,3 dichloro aniline. The intermediates 4 and 6 synthesized were then coupled under basic conditions to deliver the target compound 7.

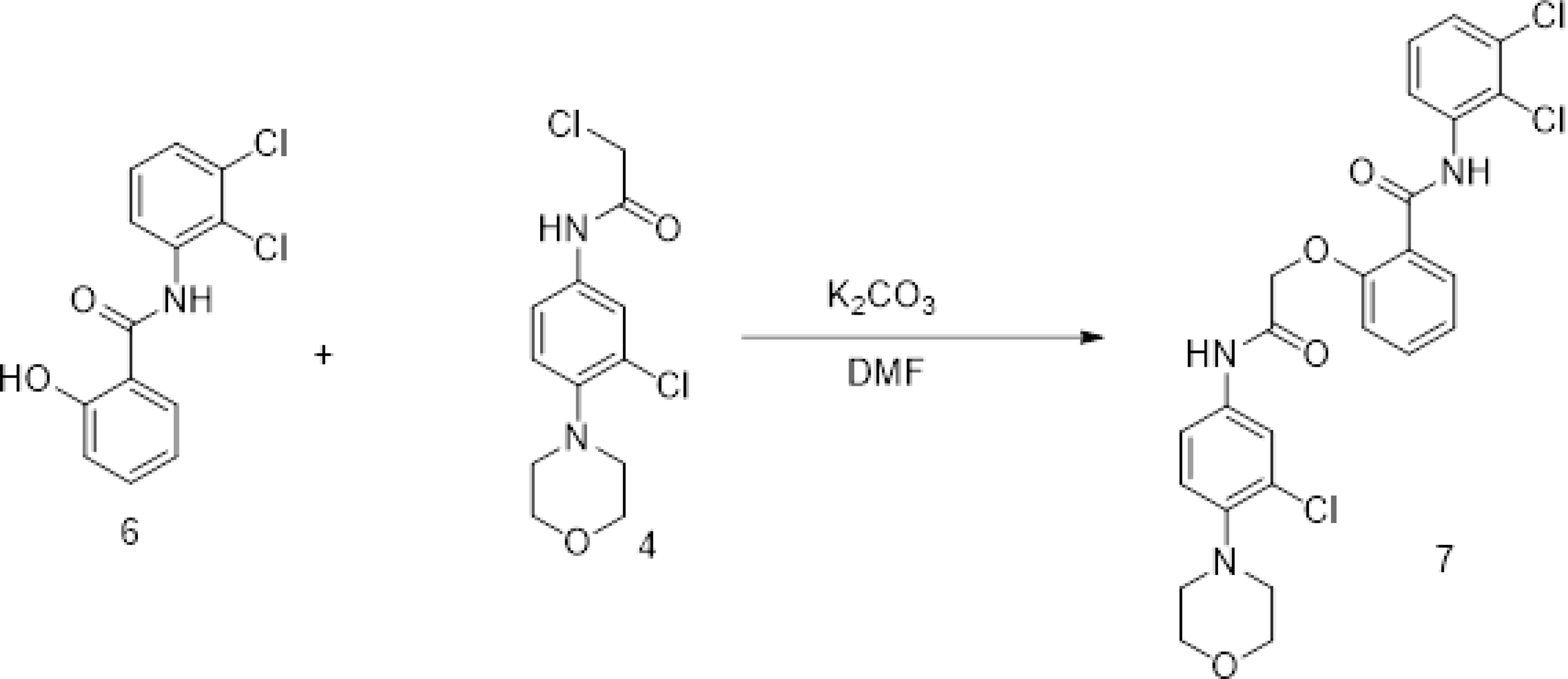

Synthesis of 2-Chloro-N-(3-chloro-4-morpholinophenyl)acetamide (4)

To a stirred solution of 3-chloro-4-morpholinoaniline (3) (1.0 g) in DCM (20 ml) was added TEA (1.5 eq.) followed by chloroacetylchloride (1.1 eq.) at 0 and 5°C. The reaction mixture was gradually warmed to RT and stirred at RT for 4h. The reaction was monitored by LCMS. After completion of reaction was quenched with ice cold water (30 mL) and extracted with DCM (2 x 30 mL). The combined organic layers were dried over sodium sulphate and concentrated over vacuo to afford crude was purified by flash chromatography (40 % EA:Hexane) to afford 2-chloro-N-(3-chloro-4-morpholinophenyl)acetamide (4) (1.2g) as brown solid. LCMS (M+H): 289.

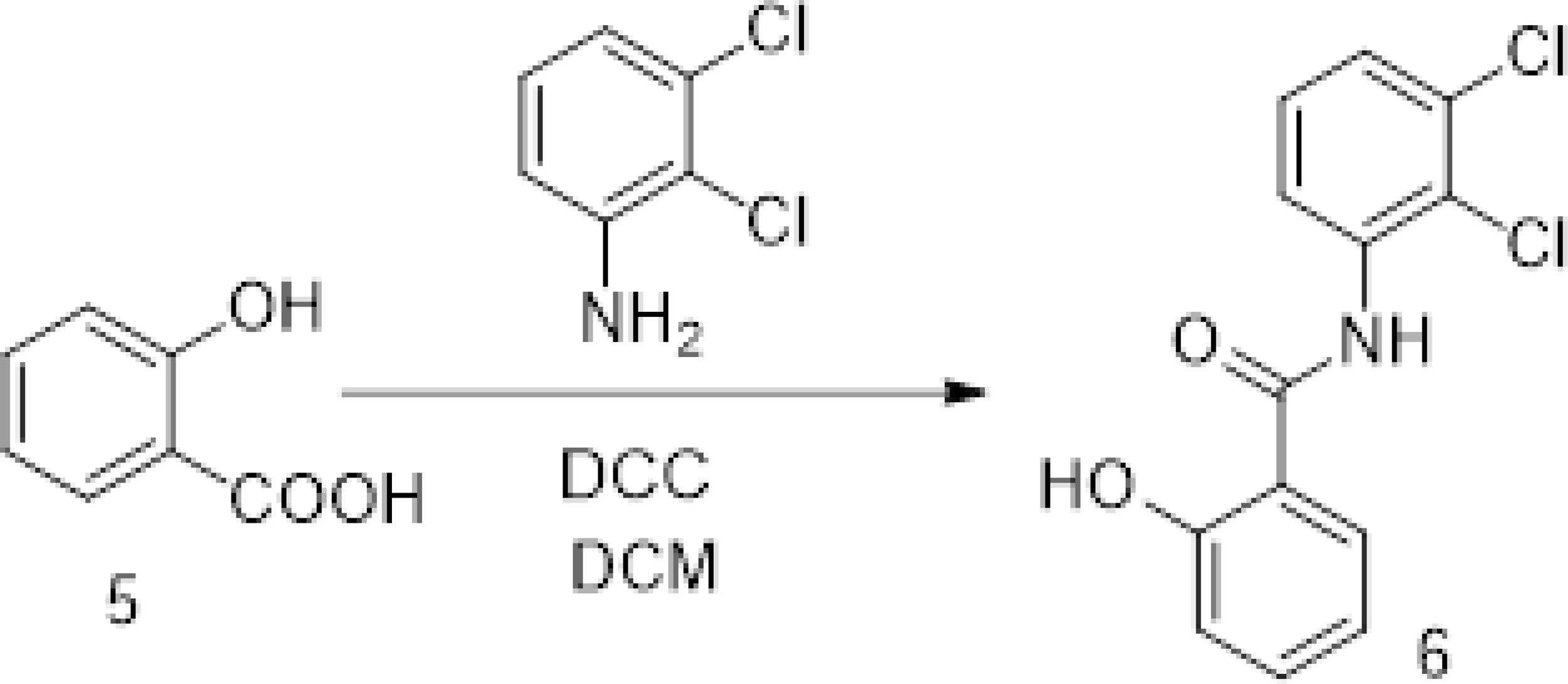

Synthesis of N-(2,3-dichlorophenyl)-2-hydroxybenzamide (6)

To a stirred solution of salicylic acid (5) (1g) and 2,3-dichloroaniline (1 eq.) in DCM (20 mL) was added (1-(3-dimethylaminopropyl)-3-ethylcarbodiimide) (1.5 eq.). The reaction mixture was heated to reflux for overnight. The reaction was monitored by LCMS. After completion of reaction the reaction mixture was concentrated under vacuo, the crude was purified by flash coloumn chromatography (30-80 % EA:Hexane) to afford N-(2,3-dichlorophenyl)-2-hydroxybenzamide (6) (500 mg) white solid. LCMS (M+H): 282.

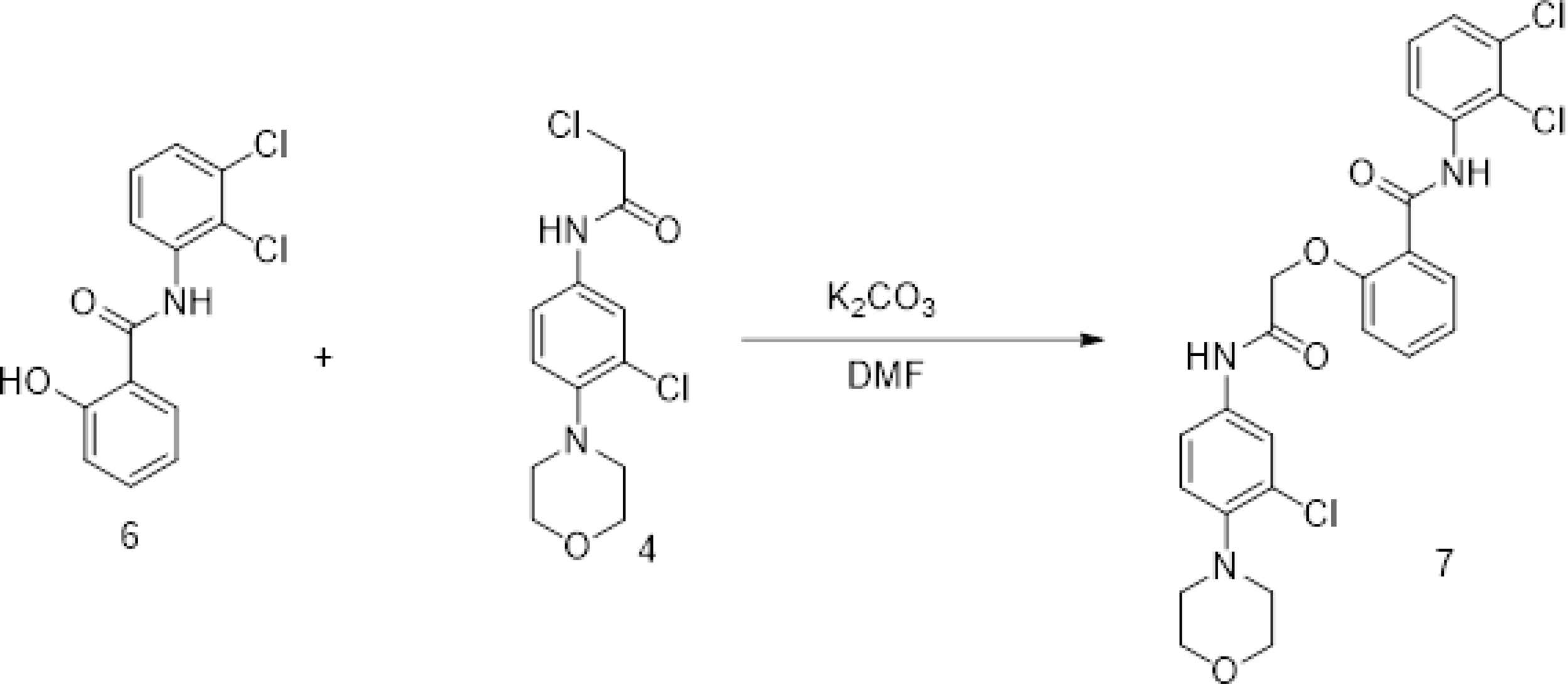

Synthesis of 2-(2-((3-Chloro-4-morpholinophenyl)amino)-2-oxoethoxy)-N-(2,3-dichlorophenyl)benzamide (7)

To a stirred solution of N-(2,3-dichlorophenyl)-2-hydroxybenzamide (6) (500 mg) in DMF (10 mL) was added anhydrous K2CO3 (2 eq.) followed by 2-chloro-N-(3-chloro-4-morpholinophenyl) acetamide (4) (1.1eq.). The reaction mixture was heated to 80°C for 16 hours. The reaction was monitored by LCMS. After completion of reaction, the reaction mixture was quenched with ice cold water (30 mL) and extracted with ethyl acetate (2 x 30 mL). The combined organic layers were dried over sodium sulphate and concentrated over vacuo to afford crude was purified by flash chromatography (3 % MeOH:CHCl3) to afford 2-(2-((3-chloro-4-morpholinophenyl)amino)-2-oxoethoxy)-N-(2,3-dichlorophenyl)benzamide (7) (400 mg) as off white solid.

LCMS (M+H): 534. 1H NMR (400 MHz, DMSO-d6) δ 10.66 (s, 1H), 10.41 (s, 1H), 8.2 (d, J = 8 Hz, 1H), 7.9 (d, J = 8 Hz, 1H), 7.7 -7.72 (m, 1H), 7.58 – 7.63 (m, 1H), 7.38 – 7.47 (m, 3H), 7.1 – 7.24 (m, 3H), 5.1 (s, 2H), 3.66 – 3.74 (m, 4H), 3.86 – 3.93 (m, 4H).

BLINK15 was synthesized from leap chem co. ltd. The synthesis of BLINK15 with smiles COc1ccc(cc1N+=O)C(=O)C[C@@]2(O)C(=O)N(c(c23)ccc(Br)c3)Cc4ccccc4 begins with 4-nitroanisole (COc1ccc(cc1N+=O)) as the starting material. First, introduce the formyl group at the para position relative to the methoxy group using a formylation reaction, such as the Vilsmeier-Haack reaction, to obtain 4-nitro-2-methoxybenzaldehyde (COc1ccc(cc1N+=O)C(=O)H). Next, perform an Aldol condensation to react the formylated product with an appropriate enolate, forming the α,β-unsaturated ketone intermediate (COc1ccc(cc1N+=O)C(=O)CH=CH2). Proceed with a Michael addition using a suitable nucleophile to introduce the side chain, preparing the molecule for further cyclization. Then, use intramolecular cyclization to form the bicyclic lactam structure, creating the C2-O-C3 bond and resulting in the fused ring system (C[C@@]2(O)C(=O)N(c(c23)ccc(Br)c3)). Introduce the bromine atom through an electrophilic aromatic substitution reaction with bromine (Br2) in the presence of a catalyst, obtaining the bromo-substituted aromatic ring.

### Cell Culture

HEK293T, HepG2, and N2A cells were cultured in DMEM with 10% FBS and 1x anti-anti solution at 37°C in a 5% CO2 atmosphere. The cells were tested negative for mycoplasma Transfection in N2A cells was performed using Lipofectamine-3000, co-transfecting with Cdk5/p35 and Cdk5/p25 plasmids to study inhibition patterns. HEK293T, HepG2, and N2A cells were maintained in DMEM -High glucose medium, supplemented with 10% FBS and 1x antibiotic-antimycotic solution, under conditions of 37°C and 5% CO2. To evaluate compound toxicity, an MTT assay was performed at varying concentrations. Six hours after cell seeding, the compounds were added in different amounts, and the cells were harvested after 24 hours for further analysis. N2A cells were transfected using Lipofectamine-3000 reagent, co-transfecting with plasmids encoding CDK5/p35 and CDK5/p25 for 12 hours, followed by inhibitor treatment to assess their effects in the cell culture system.

### Animal Model

Experiments followed the institution’s animal ethics committee guidelines. Male C57/BL mice were divided into two groups, one fed a standard diet and the other a high-fat diet for three months. All experiments were conducted in compliance with the guidelines set by the institution’s animal ethics committee. Mice were housed under standard conditions of humidity and temperature, with a 12:12-hour light-dark cycle throughout the study. Two groups of four-week-old male C57BL/6J mice were established. One group (n = 8) was fed a standard laboratory mouse chow diet (Protein: 20% kcal, Carbohydrate: 70% kcal, Fat: 10% kcal), while the other group (n = 40) was placed on a high-fat diet (Protein: 20% kcal, Carbohydrate: 20% kcal, Fat: 60% kcal) (Saha *et al*, 2023). Body weight, ITT, and GTT were assessed. Mice were treated with BLINK11 and BLINK15 inhibitors at 20 mg/kg and 40 mg/kg doses for 20 days. The study included six groups: Control, HFD, HFD_BLINK11_20, HFD_BLINK11_40, HFD_BLINK15_20, and HFD_BLINK15_40. Fig EV1 shows the flow chart of the HFD model generation and drug treatments.

### GTT And ITT assay

For the glucose tolerance test, 2g/kg of glucose was utilized, and 0.75U/kg of human insulin was employed The glucose tolerance test (GTT) and insulin tolerance test (ITT) were performed in high fat diet mice models after 20 weeks of high-fat diet feeding followed by 20 days of drug administration (Maeda *et al*, 2002). For GTT the mice were kept in a fasting period for overnight before glucose injections of 2g/kg intraperitoneally. For ITT the mice were kept in a fasting for 6 hrs prior to insulin injection of 0.75 U/kg intraperitoneally. A one-touch glucometer was used to assess the blood sugar levels at various intervals (0, 60, and 120 minutes) following intraperitoneal infusion of both glucose and insulin (Saha *et al*, 2023; Smith *et al*, 2019; Website; Benedé-Ubieto *et al*, 2020). After 20 weeks of a high-fat diet and 20 days of drug administration, the tests were carried out.

### Immunoprecipitation and Kinase Assays and immunoprecipitation

The test tube CDK5 Kinase assay was performed utilizing CDK5/p35 and CDK5/p25 complexes as kinases, following the modified procedure outlined in a previous study (Binukumar, Shukla, Amin, Bhaskar, et al., 2015), using a Promega ADP-GloTM Kinase Assay kit. The reaction was carried out with 10 μM ATP and 2 μg of histone as the substrate for CDK5 complexes. To study the inhibition patterns, 19 CDK5 inhibitors (BLINK1-19) identified through molecular docking were added to the CDK5 kinase reaction in increasing concentrations ranging from 0 to 100 nM. The IC50 (half-maximal inhibitory concentration) values of all the inhibitors were calculated based on Log(inhibitor) vs. normalized response using GraphPad software.

Cdk5 was immunoprecipitated from N2A cells and brain lysate using a Cdk5 antibody. Kinase assays were performed using a Promega ADP-Glo™ Kinase Assay kit with Cdk5/p35 and Cdk5/p25 complexes. Histone was used as a substrate to test kinase activity CDK5 was immunoprecipitated from N2a cells overexpressing CDK5/p35 and CDK5/p25, following treatment with BLINK11 and BLINK15 inhibitors at concentrations of 50nM and 100nM, using a CDK5 antibody. A kinase assay was subsequently conducted as described in Bankston et al (Bankston *et al*, 2017)(Binukumar *et al*, 2015a) Additionally, CDK5 was precipitated from mouse brain lysates treated with BLINK11 and BLINK15 inhibitors. The immunoprecipitation was performed with 100 μL of A/G beads, 2 μg of CDK5 antibody, and 150 μg of cell and brain lysates, with elution carried out using a urea elution buffer. The kinase assay was conducted using the ADP-Glo Kinase Assay Kit (Promega) with immunoprecipitated CDK5, ATP, and 1-2 μg of histone as the substrate.

### Behavioral experiments

The darkest time of the housing state was used for all behavioral studies. An hour before the experiment, the mice were acclimated in the behavior setup room. The Anymaze software (ANY-maze Video Tracking Software from Stoelting Co, 620 Wheat Lane Wood Dale, IL 60191 USA) was used for video tracking and data collection.

#### Grip strength

A grip strength meter was used to measure the forelimb grip strength in mice (Meyer *et al*, 1979). The mouse was given a chance to grab the gauge with its forelimbs before having its tail gently pulled back. Digital transducers were used to record the muscular force. Five trials were conducted on each mouse (Saha *et al*, 2023).

#### Rotarod

Rats and mice have been tested for motor coordination using the rotarod test. The animals were placed on a horizontal rod that rotates around its long axis for this test; in order to remain upright and prevent sliding off, the animal must move ahead. Every five minutes, the rod’s speed was raised, and the number of falls were counted to assess the motor coordination (Saha *et al*, 2023; Deacon, 2013).

#### Elevated plus maze

In order to examine anxiety-like behavior, phenotype transgenic and knockout mice, and screen for anxiolytic and anxiogenic medicines, this maze takes advantage of the natural fear of open, high areas in rodents (Walf & Frye, 2007; Komada *et al*, 2008). Each animal had a 10-minute test, during which we measured the entry into the open and closed arms as well as the time and distance spent in each component.

The Elevated Plus Maze (EPM) experiment is a widely utilized technique in behavioral neuroscience to evaluate anxiety-related behaviors in mice (Komada *et al*, 2008). The apparatus comprises two open arms and two closed arms arranged in a plus-shaped configuration and elevated above the ground. Mice are introduced to the center of the maze, and their movements are observed over a set period of 5 minutes (Pellow *et al*, 1985; Walf & Frye, 2007). The amount of time spent in the open arms versus the closed arms is recorded, with more time in the open arms indicating lower anxiety levels. The distance travelled in each arm were also analyzed using anymaze software. This method leverages the natural tendency of rodents to avoid open spaces and seek enclosed areas

- Y-**maze test**
- **Z-** As previously mentioned, the Y-maze test was utilized to evaluate the mouse groups’ short-term spatial memory The Y-maze test was used to assess short-term spatial memory in the mouse groups. The apparatus consists of three arms arranged in a Y-shape, typically labeled A, B, and C, with each arm having the same length and width. Mice were initially placed in one arm (usually arm A) and allowed to explore freely for 5 minutes, with one of the other arms blocked off, restricting exploration to the starting arm and the remaining open arm. After a 5-minute delay, the mouse was reintroduced to the maze with all arms accessible. The time spent in each arm and the number of entries into each arm were recorded over a 5-minute period. A higher number of entries into the previously blocked arm indicates improved spatial memory and cognitive function, suggesting the mouse recognized the novel arm. We also evaluated the mice’s spontaneous alternation rate during the Y-maze test. Spontaneous alternation reflects working memory, as it refers to the mouse’s tendency to enter a less recently visited arm. The sequence and number of arm entries were tracked, with an alternation defined as consecutive entries into all three arms without repetition (e.g., A-B-C, B-C-A) (Kraeuter *et al*, 2019; Kim *et al*, 2023; Prieur & Jadavji, 2019). During the training phase, one arm (identified as the new arm) was blocked in an indistinct, Y-shaped area that made up the maze. During the training phase, a single mouse was set free at the lower arm & given five minutes to explore the maze. The mice underwent the test phase for five minutes apiece following an hour-long intertrial break, during which time the previously blocked arm (new arm) was now available. As a result, we evaluated the novel arm’s exploration according to many criteria.

Spontaneous Alternation (%) = {(Number of alternations)/ (Total arm entries−2)} ×100

#### The Morris-water Maze (MWM) test

The Morris Water Maze (MWM) test was conducted to evaluate memory and spatial learning in mice, using visual cues placed on the walls of a water tank. For five consecutive days, the mice were trained in the water maze, with each session involving the release of the mice from various starting points and the task of locating a hidden platform within the maze. Each day, the mice underwent four trials, following the established protocol by Charles V. Vorhees (2006). The time taken (latency) for each mouse to find the platform was measured and recorded throughout the training period.

On the sixth day, a probe test was conducted to assess reference memory. The hidden platform was removed, and the mice’s behavior was observed as they navigated the maze. The time spent in the target quadrant and the time taken to reach the former platform location were recorded. This probing phase aimed to evaluate the mice’s retention of the spatial information and memory acquired during the training sessions. The Morris water maze (MWM) test evaluated memory and spatial learning in mice using visual cues in a water tank. Over five days, mice were trained to find a platform within the tank, undergoing four trials daily as per Charles V. Vorhees’ protocol. On day six, a probing experiment assessed reference memory. The platform was hidden with opaque water, and mice behavior was observed as they navigated the tank. Time spent and latency to reach the target quadrant were recorded. This phase gauged the retention of spatial information and memory acquired during training.

### PAMPA assay

Parallel Artificial Membrane Permeation Assays (PAMPA) assay was performed using a PAMPA filter plate (MAIPNTR10) coated with brain polar lipid extract (141101P) to evaluate blood brain-barrier (BBB) permeability of the compounds. The assay was performed with the help of the protocol provided by Merck Millipore on their official website and articles as cited (Website; Di *et al*, 2003). 5% of brain polar solution was used to coat the donor plate (15μl per well) by drying the plate in a fume hood. The acceptor plate was filled with 300ul of buffer (5% DMSO in PBS). The donor plate coated with brain polar solution was placed above the acceptor plate and different concentrations of Cdk5 inhibitor (100-500μM) dissolved in 5% DMSO were added to it (<u>15</u>μl per well). After 7 hrs of incubation, the absorbance was measured at broad spectrum 250-500 nm. The absorbance at 300nm was considered for further calculations as it showed an increasing pattern of ODs with an increase in the concentration of the inhibitors. For permeability, the P_app_ of the drugs was calculated.

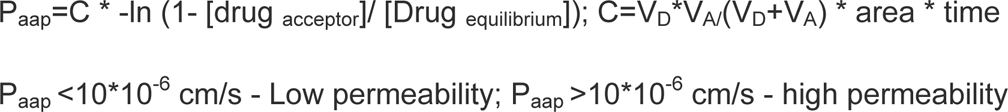

### Western Blotting

For tissue lysate preparation the mice were sacrificed following the Institution’s ethical guidelines. The hippocampus was isolated from the rest of the brain and treated in a tissue lysis solution with a protease and phosphatase inhibitor cocktail. Western blotting was performed to study the expression patterns of the protein of interest. Each well of a polyacrylamide gel was loaded with an identical amount of protein lysate (40–50 μg), which was then transferred to the nitrocellulose membrane. In 1X PBS with 0.1% (v/v) Tween 20, the main antibodies were diluted. (A list of the antibodies utilized is provided in Table EV1).

**Table EV1:**
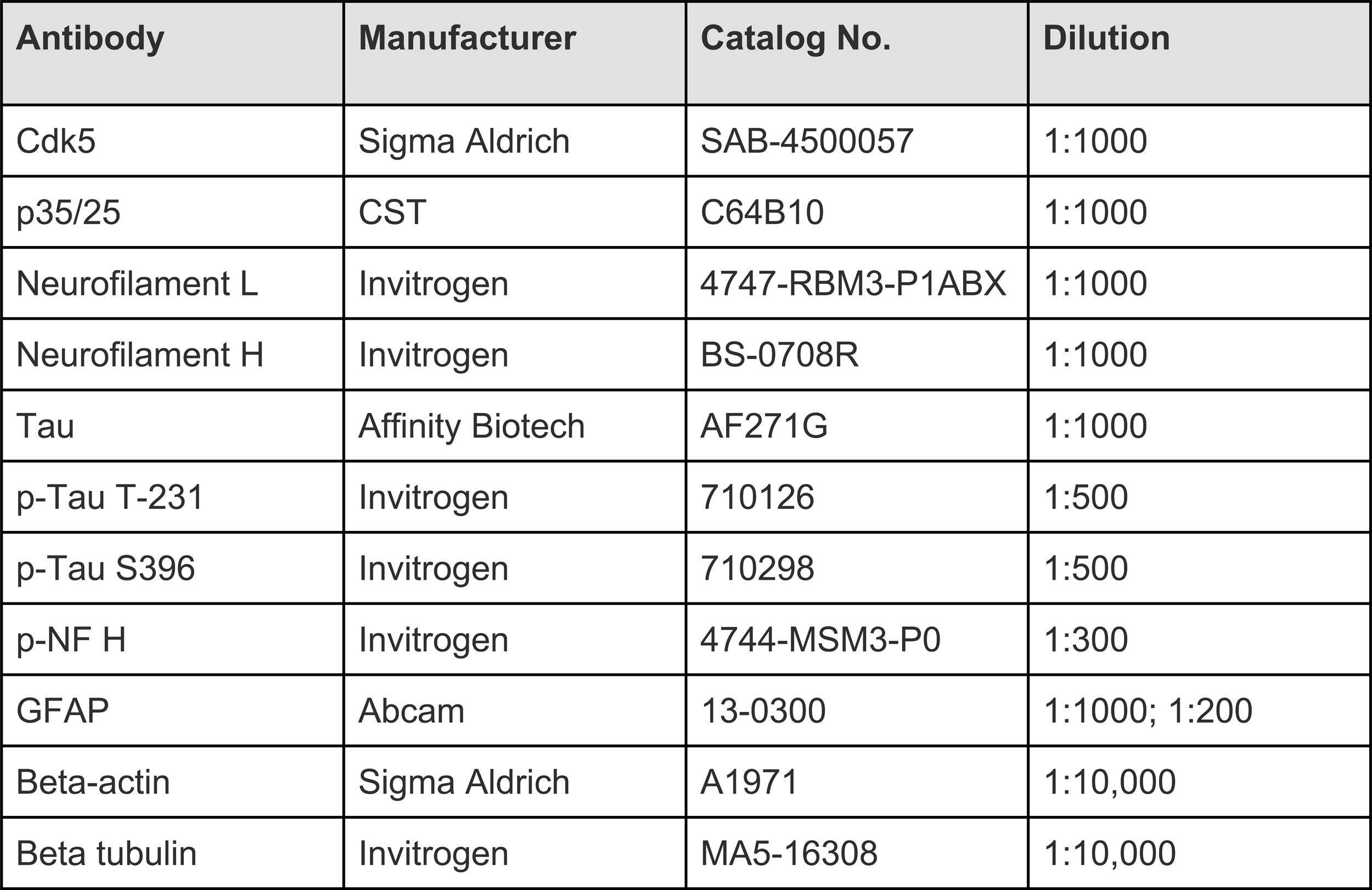
List of antibodies used in the study along with their manufacturer, catalog number, and dilutions used for experiments.

### Immunohistochemistry

The brain stored in 43% formaldehyde was processed for parafilm block preparation. The brain embedded in paraffin blocks was cut into slices with a 5 μm thickness, which were then assembled on coated glass slides. The methods for single or double immunostaining were followed while doing immunohistochemistry (Bk *et al*, 2019). The primary antibodies utilized were NF-L (1:100), Tau (1:100), p-tau S396 (1:100), p-tau T-231 (1:100), and GFAP (1:300). Utilizing secondary antibodies from Invitrogen’s Alexa fluor line that produce red (488 nm) and green (594 nm) light, immunostaining was seen. Confocal microscopy (Nikon Eclipse Ti2) was used to obtain the images of the sections from mainly the hippocampus region.

### The precise compound concentration determination in plasma and brain tissues using advanced mass spectrometry techniques

Metabolites, comprising the entire spectrum within the lysate excluding proteins but inclusive of drugs, were extracted from plasma and brain tissue homogenate samples. The extraction process involved precipitating proteins through the addition of methanol. In a concise procedure, 600 µl of methanol was introduced to 200 µl of the sample, followed by vortex mixing. The mixture was then incubated for 30 minutes in a -20℃ freezer. This method facilitated the isolation of metabolites for subsequent analysis. The solution was centrifuged at 15000 g at 4℃ for 10 minutes, and the supernatant was collected in a fresh tube and dried using a vacuum concentrator operated at room temperature. Extracted metabolites were reconstituted in 5 ml of 50% methanol with 0.1% formic acid and transferred to HPLC vials. Compounds were analyzed in these samples using a multiple reaction monitoring (MRM) method (using parameters mentioned in Table EV2) on a triple quadrupole hybrid ion trap mass spectrometer (QTRAP 6500+, SCIEX) coupled with an ExionLC UHPLC system (SCIEX). Optimized source and gas parameters were used and data acquisition was performed through Analyst 1.6.3 software in positive ion scan mode. 10 µl of metabolites suspension was loaded and resolved on an Acquity UPLC BEH C18 column (1.7 μm, 2.1 × 100 mm) using mobile phases of water with 0.1% formic acid (buffer A) and acetonitrile with 0.1% formic acid (buffer B) with a flow rate of 0.25 ml/min and 8 minutes long gradient. The gradient program was employed as follows: 2% buffer B for an initial 0.5 minutes which was ramped to 50% in the next 1.5 minutes. Buffer B was further increased to 90% in the next 1 minute and kept constant at the same concentration for the next 2 minutes. Buffer B concentration was brought to an initial 2% in the next 1 minute and the column was equilibrated at this buffer condition for 2 minutes before the next sample injection. Relative quantification was performed using MultiQuantTM software v.3.0 (SCIEX). A pooled QC from samples was used to analyze the technical variability and only metabolites with <30% coefficient of variation (%CV) were quantified.

**Table EV2:**
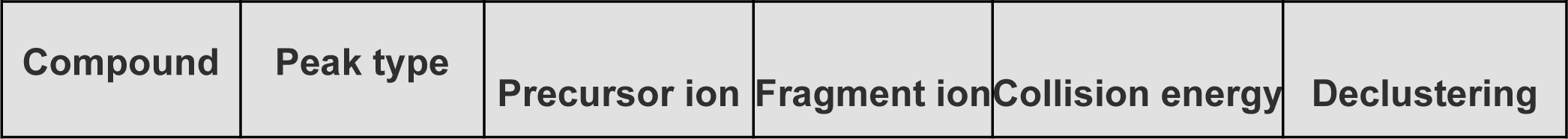
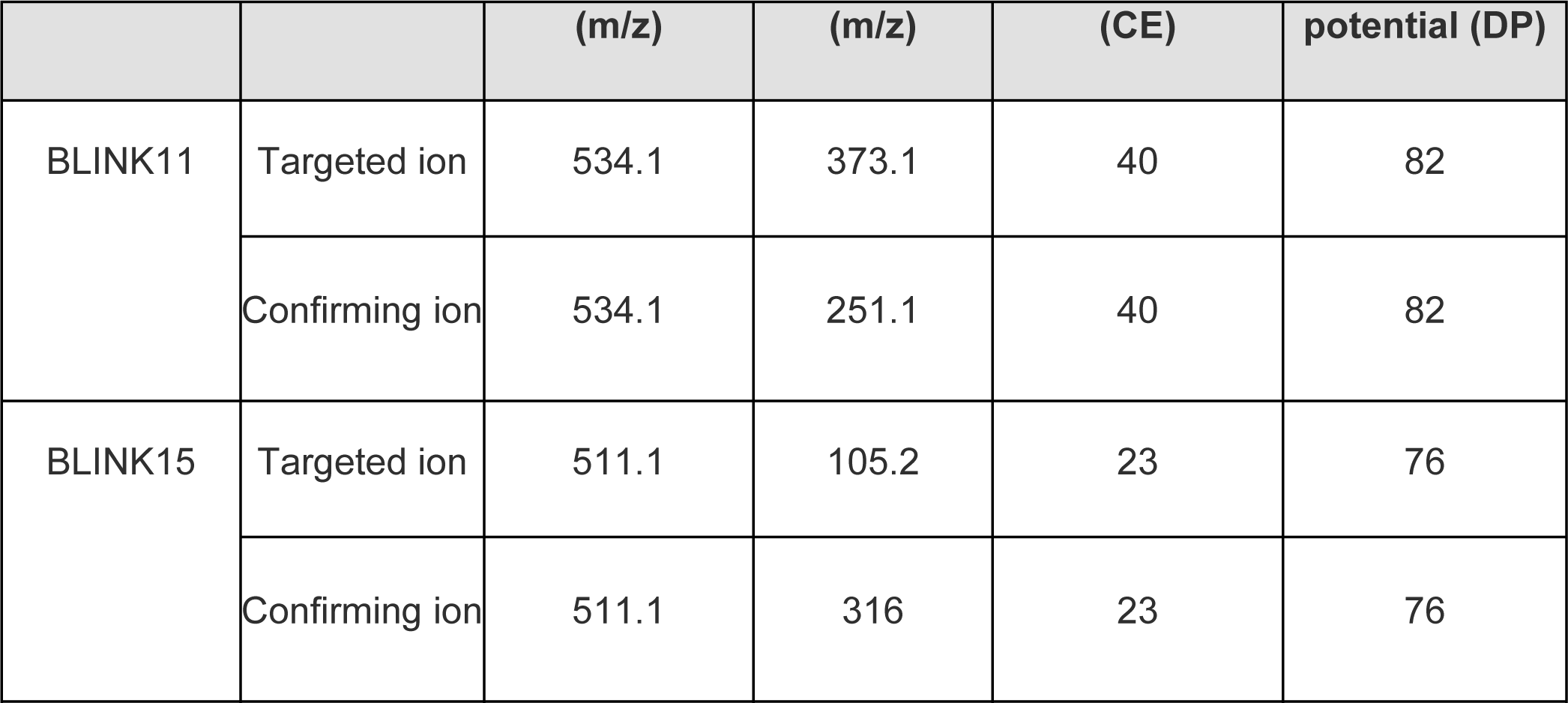
Parameters used for compound screening in the MRM method.

### 2.14 Statistical analysis

Statistical analysis was conducted using GraphPad Prism 5. Data were presented as mean ± SD. One-way ANOVA and Tukey post hoc tests determined statistical significance. IC50 values were calculated using log-transformed data. Statistical trends were indicated in figure legends, with p-values <0.05, <0.01, and <0.001 denoted by *, **, and ***, respectively. GraphPad Prism 5 was used to do statistical analysis on all datasets. The mean + SD was used to show each data set. To determine the degree of statistical importance between the animal groups, a one-way ANOVA and a Turkey post hoc test were used. The figure legends have revealed statistical trends (p-value significance). Between the specified groups, * indicates P < 0.05, **P < 0.01, and ***P < 0.001. The IC50 of inhibitors was calculated using the logarithm of x and normalized y values.

## Graphics

The graphical representation image and Fig EV1 were created using BioRender.com, and all the figures were labelled using Adobe Illustrator.

## Author contributions

BK. conceived the presented idea. S.P. carried out the in vitro and in vivo experiments. D.K.V, and R.C, performed all the molecular modeling studies. S.P., D.K.V, R.C., and B.K. wrote the manuscript. J.B. made the graphical abstract. P.S. performed the mass spectroscopy experiments. PS, SC, and SAM collaborated in synthesizing the BLINK11 compound. S.P., J.B. and B.K. contributed to the final version of the manuscript.

## Acknowledgment

This research received financial backing from CSIR India, specifically through grants OLP2301 and MLP2008, as well as support from the Science and Engineering Research Board (SERB) India, through grant GAP0168. CSIR provided senior research fellowships for SP and JB. R.C. acknowledges DBT-RA and CMNKPDF fellowships. The funders played no role in the decision to publish or in the manuscript’s creation. Special thanks go to Mukesh Kumar for his valuable assistance in performing the mice euthanization procedure during the animal experiments. We express our gratitude to the IGIB Mass Spectrometry Facility run by Shantanu Sengupta lab for their valuable assistance in quantifying the kinase inhibitors in both brain and plasma samples.

The Paper explained

## Problem

Type 2 diabetes (T2D) is linked to cognitive decline and Alzheimer’s disease (AD), both of which involve dysregulation of Cdk5 activity. Current treatments for T2D-related cognitive decline are limited, so more effective therapies are needed.

## Result

The study identified two novel Cdk5 inhibitors, BLINK11 and BLINK15, through the KINACore library. These inhibitors, particularly BLINK11, showed promising results in improving T2D symptoms (blood glucose, obesity) and reversing cognitive impairment in a high-fat diet-induced T2D model.

## Impact

Targeting Cdk5 with brain-penetrant inhibitors, like BLINK11, offers a new therapeutic approach for treating both the metabolic and cognitive impairments associated with T2D, potentially improving patient outcomes.

## Expanded View tables

**Table EV3:**
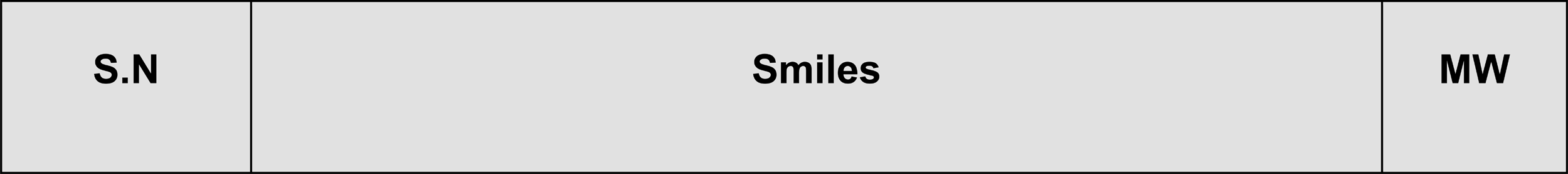
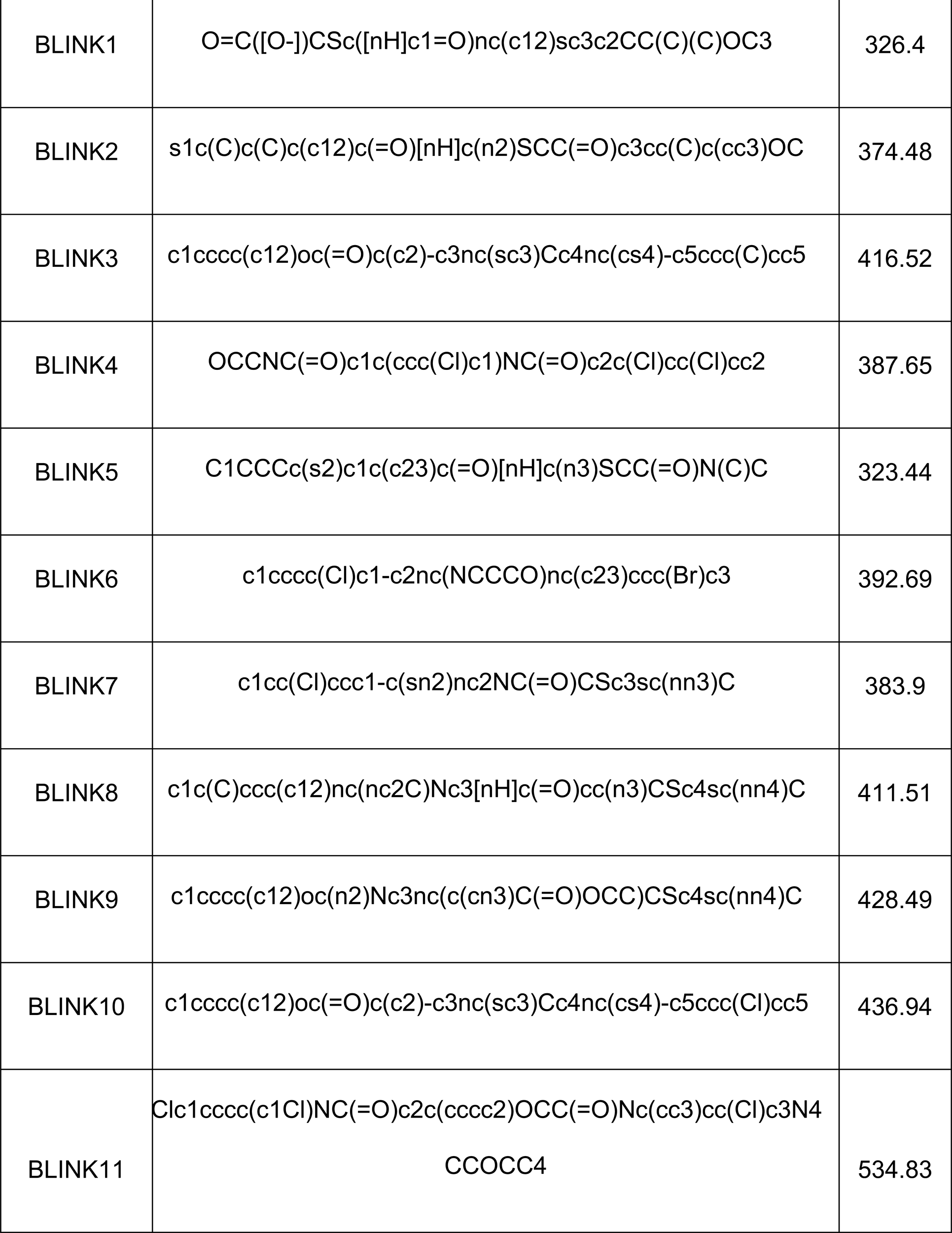
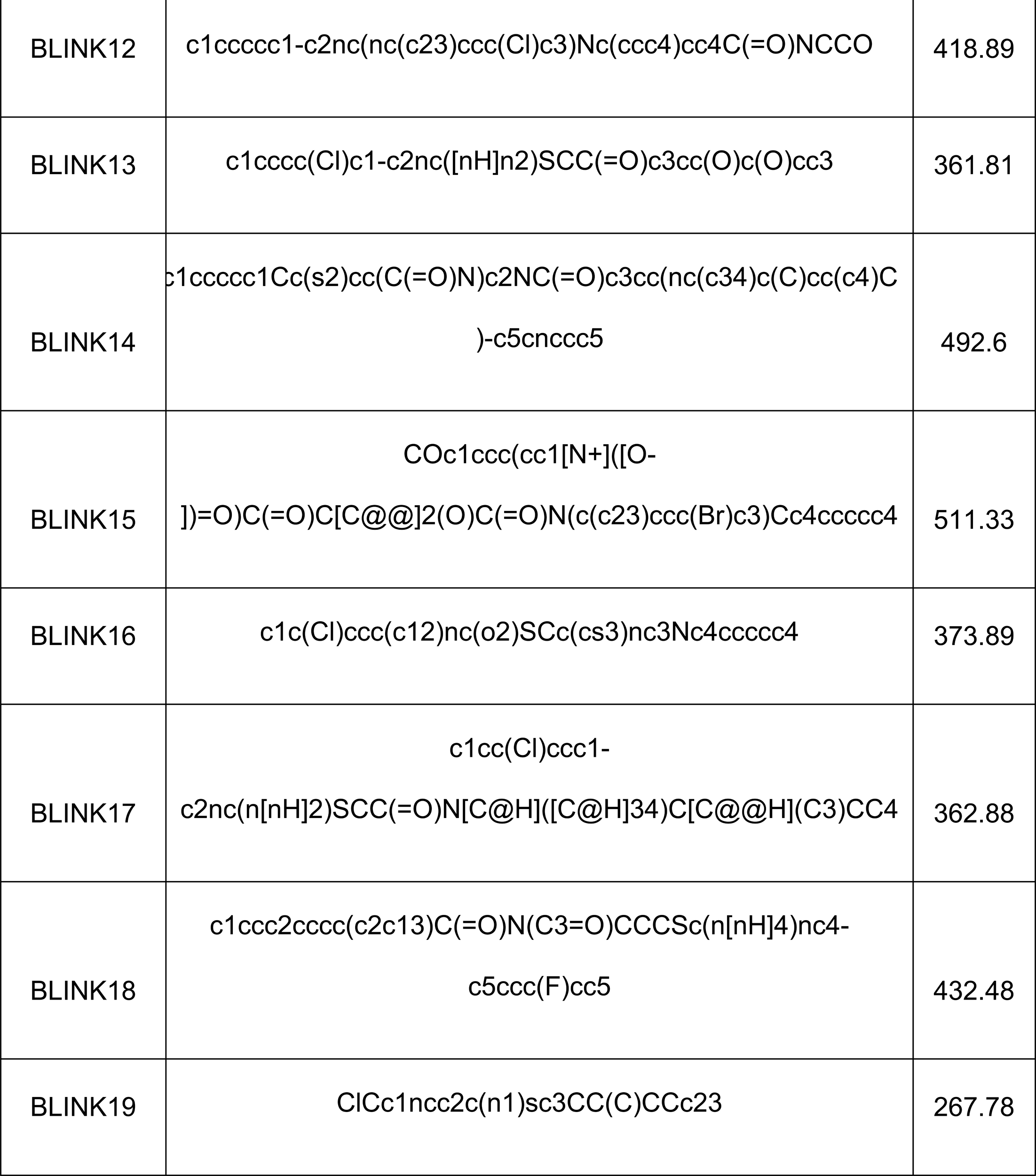
The details of the 19 selected ligands obtained from screening of KINACore libraries against Cdk5 for in vitro studies along with their corresponding molecular formula, SMILES notation, and molecular weight.

## Expanded View Figure legends

**Fig EV1:** The flowchart of HFD mice model generation and drug treatment.

**Fig EV2:** Structure of all the 19 compounds (BLINK1-19) selected by in silico approach

**Fig EV3:** RMSD plot determined for Cdk5-BLINK complexes (during 100ns simulation) specifically BLINK1-19, which were sequentially depicted from A to Q. In each complex, Cdk5 is denoted by black lines, while the corresponding ligands are illustrated with red lines. The RMSD plots for the interactions of BLINK11 and BLINK15 with Cdk5 are individually displayed in Figures 3F and 3G, respectively.

**Fig EV4: PAMPA assay** of BLINK11 and BLINK15 compound (A-B). **Evaluation of non-specific kinase inhibition by BLINK15 and BLINK11.** The graph shows the inhibition pattern of (C) Cdk1/CylY kinase activity (D) Cdk2/CylA2 and (E) Cdk9/CylT by BLINK11 and BLINK15 with increasing concentration (0-100nM). (F) Shows IC50 of all the kinases for both BLINK11 and BLINK15 inhibitors.

